# Identifying Transcriptomic Correlates of Histology using Deep Learning

**DOI:** 10.1101/2020.08.07.241331

**Authors:** Liviu Badea, Emil Stănescu

## Abstract

Linking phenotypes to specific gene expression profiles is an extremely important problem in biology, which has been approached mainly by correlation methods or, more fundamentally, by studying the effects of gene perturbations. However, genome-wide perturbations involve extensive experimental efforts, which may be prohibitive for certain organisms. On the other hand, the characterization of the various phenotypes frequently requires an expert’s subjective interpretation, such as a histopathologist’s description of tissue slide images in terms of complex visual features (e.g. ‘acinar structures’). In this paper, we use Deep Learning to eliminate the inherent subjective nature of these visual histological features and link them to genomic data, thus establishing a more precisely quantifiable correlation between transcriptomes and phenotypes. Using a dataset of whole slide images with matching gene expression data from 39 normal tissue types, we first developed a Deep Learning *tissue classifier* with an accuracy of 94%. Then we searched for *genes whose expression correlates with features inferred by the classifier* and demonstrate that Deep Learning can automatically derive visual (phenotypical) features that are well correlated with the transcriptome and therefore *biologically interpretable*. As we are particularly concerned with *interpretability* and *explainability* of the inferred histological models, we also develop *visualizations* of the inferred features and compare them with gene expression patterns determined by immunohistochemistry. This can be viewed as a first step toward bridging the gap between the level of genes and the cellular organization of tissues.

## Introduction

Histological images have been used for biological research and clinical practice already since the 19^th^ century and are still employed in standard clinical practice for many diseases, such as various cancer types. On the other hand, the last few decades have witnessed an exponential increase in sophisticated genomic approaches, which allow the dissection of various biological phenomena at unprecedented molecular scales. But although some of these omic approaches have entered clinical practice, their potential has been somewhat tempered by the heterogeneity of individuals at the genomic and molecular scales. For example, omic-based predictors of cancer evolution and treatment response have been developed, but their clinical use is still limited [Xin, 2017] and they have not yet replaced the century old practice of histopathology. So, despite the tremendous recent advances in genomics, histopathological images are still often the basis of the most accurate oncological diagnoses, information about the microscopic structure of tissues being lost in genomic data.

Thus, since histopathology and genomic approaches have their own largely non-overlapping strengths and weaknesses, a combination of the two is expected to lead to an improvement in the state of the art. Of course, superficial combinations could be easily envisioned, for example by constructing separate diagnosis modules based on histology and omics respectively, and then combining their predictions. Or, alternatively, we could regard both histopathological images and genomic data as features to be used jointly by a machine learning module [Mobadersany, 2018].

However, for a more in-depth integration of histology and genomics, a better understanding of the relationship between genes, their expression and histological phenotypes is necessary. Linking phenotypes to specific gene expression profiles is an extremely important problem in biology, as the transcriptomes are assumed to play a causal role in the development of the observed phenotype. The definitive assessment of this causal role requires studying the effects of gene perturbations. However, such genome-wide perturbations involve extensive and complex experimental efforts, which may be prohibitive for certain model organisms.

In this paper we try to determine whether there is a link between the *expression of specific genes* and *specific visual features* apparent in histological images. Instead of observing the effects of gene perturbations, we use representation learning based on deep convolutional neural networks [Goodfellow, 2014] to automatically infer a large set of visual features, which we correlate with gene expression profiles.

While gene expression values are easily quantifiable using current genomic technologies, determining and especially quantifying visual histological features has traditionally relied on subjective evaluations by experienced histopathologists. However, this represents a serious bottleneck, as the number of qualified experts is limited and often there is little consensus between different histologists analyzing the same sample [Baak, 1982]. Therefore, we employ a more objective visual feature extraction method, based on deep convolutional neural networks. Such more objective visual features are much more precisely quantifiable than any subjective features employed by human histopathologists. Therefore, although this correlational approach cannot fully replace perturbational studies of gene-phenotype causation, it is experimentally much easier and at the same time much more precise, due to the well-defined nature of the visual features employed.

In our study, we concentrate on gene expression rather than other multi-omic modalities, since the transcriptional state of a cell frequently seems to be one of the most informative modalities [Yuan, 2014].

There are numerous technical issues and challenges involved in implementing the above-mentioned research.

The advent of *digital* histopathology [Aeffner, 2019] has enabled the large scale application of automated computer vision algorithms for analysing histopathological samples. Traditionally, the interpretation of histopathological images required experienced histopathologists as well as a fairly long analysis time. Replicating this on a computer was only made possible by recent breakthroughs in Deep Learning systems, which have achieved super-human performance in image recognition tasks on natural scenes, as in the ImageNet Large Scale Visual Recognition Competition (ILSVRC) [Krizhevsky, 2012; Russakovsky, 2015]. However, directly transferring these results to digital histopathology is hampered by the much larger dimensions of histological whole slide images (WSI), which must be segmented into smaller image patches (or “tiles”) to make them fit in the GPU memory of existing Deep Learning systems. Moreover, since there are far fewer annotated WSI than natural images and since annotations are associated with the whole slide image rather than the individual image tiles, conventional supervised Deep Learning algorithms cannot always be directly applied to WSI tiles. This is particularly important whenever the structures of interest are rare and thus do not occur in all tiles. For example, tumor cells may not be present in all tiles of a cancer tissue sample. Therefore, we concentrate in this paper on normal tissue samples, which are more homogeneous at not too high magnifications and for which the above-mentioned problem is not as acute as in the case of cancer samples.

Only a few large-scale databases of WSI images are currently available, and even fewer with paired genomic data for the same samples. The Cancer Genome Atlas (TCGA) [Weinstein, 2013] has made available a huge database of genomic modifications in a large number of cancer types, together with over 10,000 WSI. The Camelyon16 and 17 challenges [Litjens, 2018] aimed at evaluating new and existing algorithms for automated detection of metastases in whole-slide images of hematoxylin and eosin stained lymph node tissue sections from breast cancer patients. The associated dataset includes 1,000 slides from 200 patients, whereas the Genotype-Tissue Expression (GTEx) [Lonsdale, 2013] contains over 20,000 WSI from various normal tissues.

Due to the paucity of large and well annotated WSI datasets for many tasks of interest (such as cancer prognosis), some research groups employed *transfer learning* by reusing the feature layers of Deep Learning architectures trained on ordinary image datasets, such as ImageNet. Although histopathological images look very different from natural scenes, they share basic features and structures such as edges, curved contours, etc., which may have been well captured by training the network on natural images. Nevertheless, it is to be expected that training networks on genuine WSI will outperform the ones trained on natural scenes [Coudray, 2018].

The most active research area involving digital histopathology image analysis is computer assisted diagnosis, for improving and speeding-up human diagnosis. Since the errors made by Machine Learning systems typically differ from those made by human pathologists [Wang, 2016], classification accuracies may be significantly improved by *assisting* humans with such automated systems. Moreover, reproducibility and speed of diagnosis will be significantly enhanced, allowing more standardized and prompt treatment decisions. Other supervised tasks applied to histopathological images involve [Komura, 2018]: detection and segmentation of regions of interest, such as automatically determining the tumour regions in WSI [Spanhol, 2016], scoring of immunostaining [Sheikhzadeh, 2016], cancer staging [Wang, 2016], mitosis detection [Shah, 2017], gland segmentation [Chen, 2016], or detection and quantification of vascular invasion [Caie, 2014].

Machine Learning has also been used for discovering new clinical-pathological relationships by correlating histo-morphological features of cancers with their clinical evolution, by enabling analysis of huge amounts of data. Thus, relationships between morphological features and somatic mutations [Coudray, 2018; Chen, 2020; Schaumberg, 2018], as well as correlations with prognosis have been tentatively addressed. For example, [Yu, 2016] developed a prognosis predictor for lung cancer using a set of predefined image features. Still, this research field is only at the beginning, awaiting extensive validation and clinical application.

From a technical point of view, the number of *Deep Learning architectures*, systems and approaches used or usable in digital histopathology is bewildering. Such architectures include supervised systems such as Convolutional Neural Networks (CNN), or Recurrent Neural Networks (RNN), in particular Long Short Term Memory (LSTM) [Hochreiter, 1997] and Gated Recurrent Units [Cho, 2014]. More sophisticated architectures, such as the U-net [Ronneberger, 2015], multi-stream and multi-scale architectures [Kamnitsas, 2017] have also been developed to deal with the specificities of the domain. Among unsupervised architectures we could mention Deep Auto-Encoders (AEs) and Stacked Auto-Encoders (SAEs), Generative Adversarial Networks [Goodfellow, 2014; Gadermayr, 2018], etc.

Currently, CNNs are the most used architectures in medical image analysis. Several different *CNN architectures*, such as AlexNet [Krizhevsky, 2012], VGG [Simonyan, 2014], ResNet [He, 2016], Inception v3 [Szegedy, 2016], etc. have proved popular for medical applications, although there is currently no consensus on which performs best. Many open source Deep Learning *systems* have appeared since 2012. The most used ones are *TensorFlow* (Google [Abadi, 2016]), *Keras* [Chollet, 2015] (high level API working on top of TensorFlow, CNTK, or Theano) and *PyTorch* [Paszke, 2019].

In this paper we look for gene expression-phenotype correlations for normal tissues from the Genotype-Tissue Expression (GTEx) project, the largest publicly available dataset containing paired histology images and gene expression data. The phenotypes consist in visual histological features automatically derived by various convolutional neural network architectures implemented in PyTorch and trained, validated and tested on 1,670 whole slide images at 10x magnification divided into 579,488 tiles. A supervised architecture was chosen, as the inferred visual features should be able to discriminate between the various tissues of interest instead of just capturing the components with the largest variability in the images.

## Materials and methods

Our study of the correlations between visual histological features and gene expression involves several stages:

1. Automated *feature discovery* using representation learning based on a supervised Convolutional Neural Network (CNN) trained to classify histological images of various normal tissues; validation and testing of the tissue classifier on completely independent samples.
2. *Quantification* of the inferred histological features in the various layers of the network and their *correlation with paired gene expression data* in an independent dataset.
3. *Visualization of features* found correlated to specific gene expression profiles. Two different visualization methods are used: *guided backpropagation* on specific input images, as well as input-independent generation of *synthetic images* that maximize these features.

The following sections describe these analysis steps in more detail (see also Fig 1).

**Fig 1.**
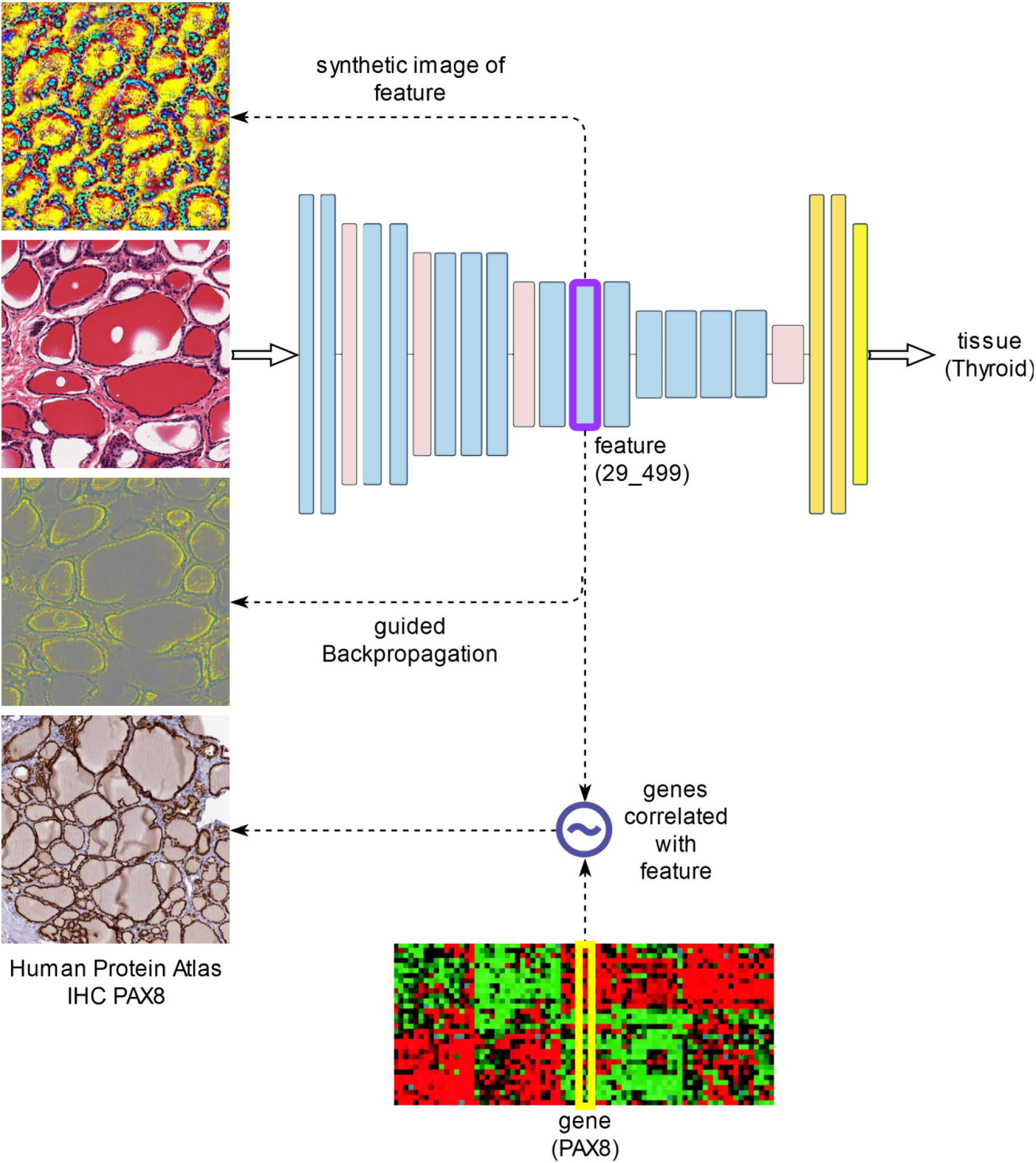
The main analysis steps.

### Developing a classifier of histological images

First, we developed a normal tissue classifier based on histological whole slide images stained with hematoxylin and eosin (H&E). Such a tissue classifier is also useful in its own right, since it duplicates the tasks performed by skilled histologists, whose expertise took years or even decades to perfect. Such a task could not even be reliably performed by computer before the recent advent of Deep Learning. But although Deep Learning has been able to surpass humans in classifying objects from natural images (as in the ImageNet competition [Russakovsky, 2015]), transferring these results to digital histopathology is far from trivial [Komura, 2018]. This is mainly because histopathology images are far larger (up to tens of billions of pixels) and labelled data much sparser than for natural images used in other visual recognition tasks, such as ImageNet. Whole slide images (WSI) thus need to be divided in tiles of sizes small enough (typically 224×224, or 512×512 pixels) to allow Deep Learning mini-batches to fit in the memory of existing GPU boards. Moreover, since there are far fewer annotated WSI than natural images and since annotations are associated with the whole slide image rather than the individual image tiles, conventional supervised Deep Learning algorithms cannot always be directly applied to WSI tiles. This is particularly important whenever the structures of interest are rare and thus do not occur in all tiles. (For example, tumor cells may not be present in all tiles of a cancer tissue sample.) Thus, the main strength of Deep Learning in image recognition, namely the huge amounts of labelled data, is not available in the case of histopathological WSI. More sophisticated algorithms based on multiple instance learning [Xu, 2017] or semi-supervised learning are needed, but haven’t been thoroughly investigated in this domain. In multiple instance learning, labels are associated to *bags* of instances (i.e. to the whole slide image viewed as a bag of tiles), rather than individual instances, making the learning problem much more difficult. In contrast, semi-supervised learning employs both labelled and unlabelled data, the latter for obtaining better estimates of the true data distribution.

Therefore, we concentrate in this paper on *normal tissue* samples, which are more homogeneous at not too high magnifications and for which the above-mentioned problem is not as acute as in the case of cancer samples. In the following, we used data from the *Genotype-Tissue Expression* (GTEx) project, the largest publicly available dataset containing paired histology images and gene expression data.

### The GTEx dataset

GTEx is a publicly available resource for data on tissue-specific gene expression and regulation [GTEx]. Samples were collected from over 50 normal tissues of nearly 1,000 individuals, for which data on whole genome or whole exome sequencing, gene expression (RNA-Seq), as well as histological images are available.

In our application, we searched for subjects for which both histological images and gene expression data (RNA-Seq) are available. We selected from these 1,670 histological images from 39 normal tissues, with 1,778 associated gene expression samples (for 108 of the histological images there were duplicate gene expression samples). The 1,670 histological images were divided into three data sets:

- *training set* (1,006 images, representing approximately 60% of the images),
- *validation set* (330 images, approximately 20% of images),
- *test set* (334 images, approximately 20% of images).

Figures 2A and 2B show such a histological whole slide image (WSI) and a detail of a thyroid gland sample.

**Fig 2.**
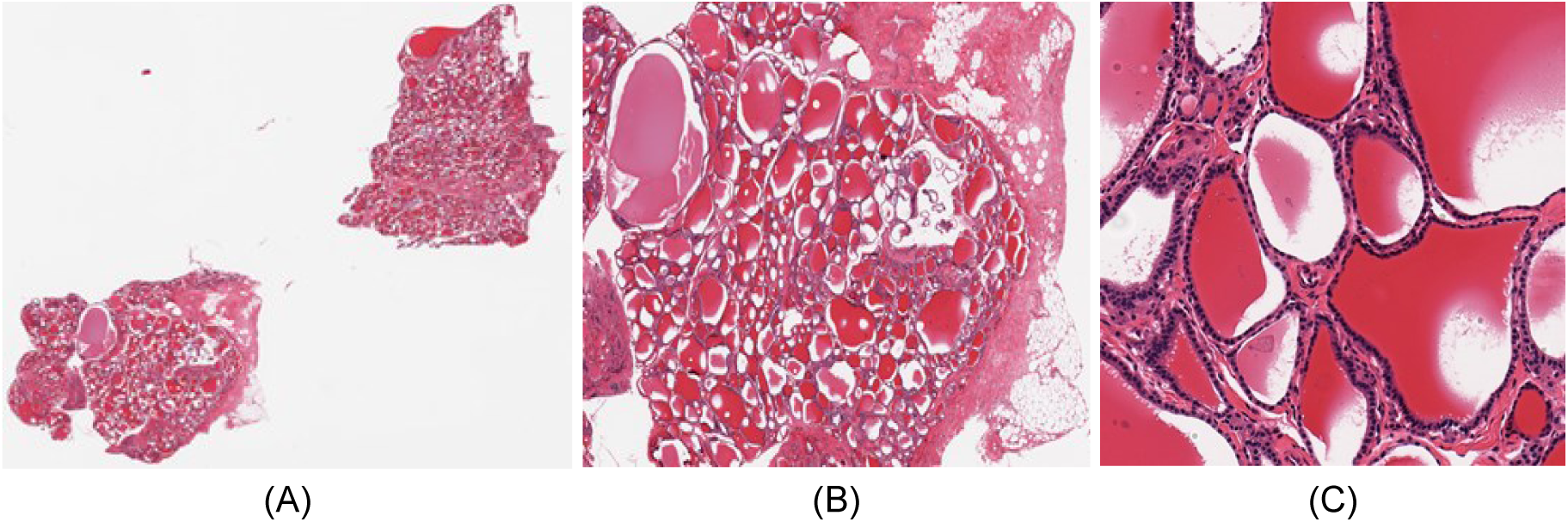
Whole slide image of thyroid sample GTEX-11NSD-0126. (A) Whole slide. (B) Detail. (C) Tile of size 512×512.

Another very important aspect specific to WSI is the optimal level of *magnification* to be fed as input to the image classification algorithms. As tissues are composed of discrete cells, critical information regarding cell shape is best captured at high resolutions, whereas more complex structures, involving many cells, are better visible at lower resolutions. The maximum image acquisition resolution in GTEx SVS files corresponds to a magnification of 20x (0.4942 mpp - microns per pixel). In this paper, we chose an intermediate magnification level of 10x (0.9884 mpp), which represents a reasonable compromise between capturing enough cellular details, and allowing a field of view of the image tiles large enough to include complex cellular structures. The whole slide images were divided into tiles of size 512×512 pixels (506μm x 506μm), which were subsequently rescaled to 224×224, the standard input dimension for the majority of convolutional neural network architectures. Image tiles with less than 50% tissue, as well as those with a shape factor other than 1 (with a non-square shape, originating from WSI edges) were removed so as not to distort the training process, and the remaining tiles were labelled with the identifier of the tissue of origin of the whole image. A 512×512 image tile of the WSI from Fig 2A can be seen in Fig 2C.

The image tiles dataset contains 579,488 such tiles (occupying 72GB of disk space) and was further split into distinct training, validation and test datasets as follows:

- 349,340 tiles in the training set (taking up 44GB of disk space),
- 114,123 tiles in the validation set (14GB),
- 116,025 tiles in the test set (15GB).

S1 Table shows the numbers of images and respectively image tiles in the three datasets (training, validation and test) for each of the 39 tissue types. By comparison, the ImageNet dataset (from the ILSVRC-2012 competition) had 1.2 million images and 1,000 classes (compared to 39 classes in the GTEx dataset).

Since to the best of our knowledge, we could not find, for validation purposes, other comparable datasets with paired histological images and gene expression data, we did not perform a color normalization optimized to the GTEx dataset, as it might not extrapolate well to other datasets with different color biases. Therefore, we used the standard color normalization employed in the ImageNet dataset. Such a normalization may be slightly suboptimal, but this is expected to have little impact on the classification results, due to the complexity of the CNNs employed.

### Training and validation of Deep Learning tissue classifiers

We trained and validated several different convolutional neural network architectures on the image tile dataset described above (Table 1).

**Table 1.**
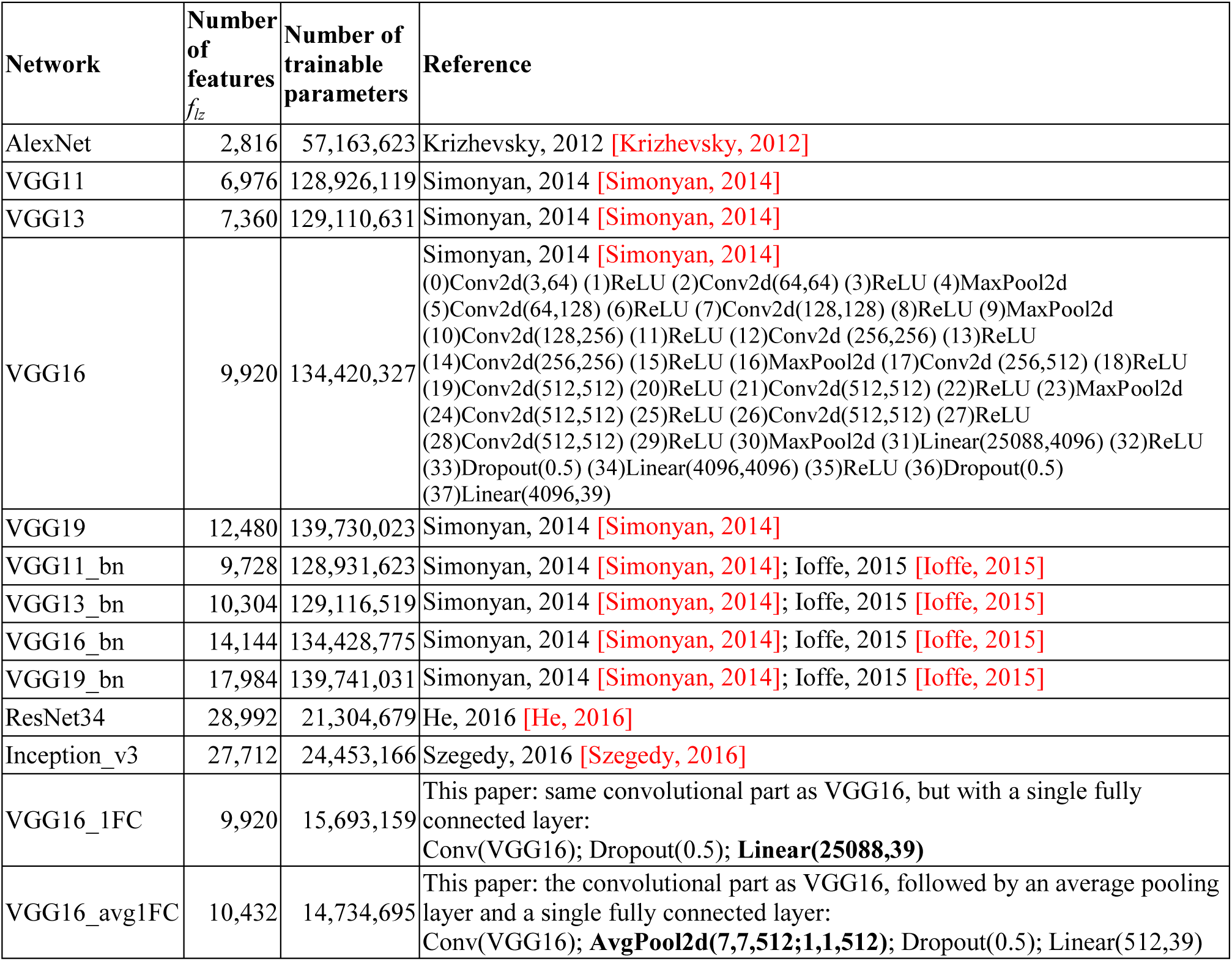
Convolutional neural network architectures used in this paper. For ResNet34, the features considered involve only layers that are not “skipped over”, while for Inception_v3 only elementary layers with non-negative outputs (ReLU, max/avg-pooling) that are not on the auxiliary branch.

The VGG16_1FC model is a modification of the VGG16 model, in which the final three fully connected layers have been replaced by a single fully connected layer, preceded by a dropout layer. This not only significantly reduces the very large number of parameters of the VGG16 architecture (from 134,420,327 to 15,693,159), but also attempts to obtain more intuitive representations on the final convolutional layers.

Since the last convolutional layer of VGG16 is of the form 7×7 x 512 channels and since location is completely irrelevant in tissue classification, we also extended the convolutional part of VGG16 by an average pooling layer (over the 7×7 spatial dimensions), followed as in VGG16_1FC by a single fully connected layer. We refer to this new architecture as VGG16_avg1FC.

Architectures with the suffix “bn” use “batch normalization” [Ioffe, 2015].

Note that in the case of ResNet34, the internal layers of the residual blocks do not make up complete representations, since their outputs are *added* to the skip connections. Therefore, the ResNet features considered here involve only layers that are not “skipped over”.

On the other hand, the Inception_v3 architecture contains many *redundant* representations, since it combines by concatenation individual multi*-*scale filter outputs (the individual filter outputs are repeated in the concatenated layer). To avoid this representational redundancy, we only consider as features the *elementary* layers with non-negative outputs (ReLU, max- or average pooling) that are not on the auxiliary branch.

We report, in the following, the accuracies of the classifiers on both *validation* and *test* datasets, after training for 90 epochs on the *training* dataset, with the initial learning rate of 0.01 and reducing it by a factor of 10 every 30 epochs. The mini-batch sizes were chosen according to the available GPU memory (11GB): 256 for AlexNet and 40 for the other models, respectively.

Note that although the validation dataset has not been used for model *construction*, it has been employed for model *selection* (corresponding to the training epoch with the best model accuracy on the validation data). Therefore, we use a completely independent *test* dataset for model evaluation.

The models were implemented in Pytorch 1.1.0 (https://pytorch.org/) [Paszke, 2019], based on the torchvision models library (https://github.com/pytorch/vision/tree/master/torchvision/models).

As the histological images contain multiple tiles, we also constructed a *tissue classifier for whole slide images (WSI)* by applying the tile classifier on all tiles of the given WSI and aggregating the corresponding predictions using the majority vote.

### Correlating histological features with gene expression data

From a biological point of view, the visual characteristics of a histological image, its phenotype, is determined by the gene expression profiles of the cells making up the tissue. But the way in which transcriptomes determine phenotypes was practically impossible to determine automatically until the advent of machine learning methods for learning representations based on Deep Learning, mainly because most histological phenotypes were hard to define precisely and especially difficult to quantify automatically.

The internal representations constructed by a convolutional neural network can be viewed as visual features detected by the network in a given input image. These visual features, making up the phenotype, can be correlated to paired gene expression data to determine the most significant gene-phenotype relationships. However, to compute these correlations, we need a precise *quantification* of the visual features.

The *quantification of the inferred visual features* for a given input image is non-trivial, as it needs to be *location invariant* – the identity of a given tissue should not depend on spatial location. Although the activations *Y*_*lzxy*_(*X*) of neurons (*x,y,z*) in layer *l* for input image *X* obviously depend on the location (*x,y*), we can construct location invariant aggregates of these values of the form:

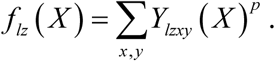

We have experimented with various values of *p*, but in the following we show the simplest version, *p*=1. This is equivalent to computing the spatial averages of neuron values in a given layer *l* and channel *z*. Thus, we employ *spatially invariant features* of the form *f*_*lz*_, which are quantified by forward propagation of histological images *X* using the formula above.

Since histological whole slide images *W* were divided into multiple tiles *X*, we also sum over these tiles to obtain the feature value for *W*:

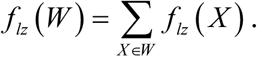

After quantification of features *f*_*lz*_(*W*_*i*_) over all samples *i*, we compute pairwise gene-feature Pearson correlations *r*(*g,f*_*lz*_), where *g*_*i*_ are the gene expression values of gene *g* in the samples *i* (for each sample *i*, we have simultaneous histological whole slide image *W*_*i*_ and gene expression data *g*_*i*_ for virtually all human genes *g*).

To avoid any potential data leakage, we compute gene-feature correlations on the *test dataset*, which is completely independent from the datasets used for model construction and respectively selection (*training* and respectively *validation* datasets).

Note that since we are in a ‘small sample’ setting (the number of variables greatly exceeds the number of samples), we performed a univariate analysis rather than a multivariate one, since multivariate regression is typically not recommended for small samples. For example, in the case of the VGG16 architecture, we have 9,920 features *f*_*lz*_, 56,202 genes (totaling 557,523,840 gene-feature pairs) and just 334 samples in the test dataset.

The selection of significantly correlated gene-feature pairs depends on the correlation-and gene expression thresholds used. Genes with low expression in all tissues under consideration should be excluded, but determining an appropriate threshold is non-trivial, since normal gene expression ranges over several orders of magnitude. (For example, certain transcription factors function at low concentrations.)

We compared the different CNN architectures from Table 1 in terms of the numbers of significant gene-feature pairs (as well as unique genes, features and respectively tissues involved in these relations) using fixed correlation-(*R*_*T*_=0.8) and *log*_*2*_ expression thresholds (*E*_*T*_=10). More precisely, the expression threshold *E*_*T*_ constrains the gene expression value *E* as follows: *log*_*2*_(1+*E*) ≤ *E*_*T*_.

We also assessed the significance of gene-feature correlations using permutation tests (with *N*=1,000 permutations).

Since one may expect the final layers of the CNN to be better correlated with the training classes (i.e. the tissue types), we studied the *layer* distribution of features with significant gene correlations.

We also studied the reproducibility of gene-feature correlations w.r.t. the dataset used. More precisely, we determined the Pearson correlation coefficient between the (Fisher transformed) gene-feature correlations computed w.r.t. the validation and respectively test datasets.

### Visualization of histological features

The issues of *interpretability* and *explainability* of neural network models is essential in many applications. Deep learning based histological image classifiers can be trained to achieve impressive performance in days, while a human histopathologist needs years, even decades, to achieve similar performance. However, despite their high classification accuracies, neural networks remain opaque about the precise way in which they manage to achieve these classifications. Of course, this is done based on the visual characteristics automatically inferred by the network, but the details of this process cannot be easily explained to a human, who might want to check not just the end result, but also the reasoning behind it. This is especially important in clinical applications, where *explainable* reasoning is critical for the adoption of the technology by clinicians, who need explanations to integrate the system’s findings in their global assessment of the patient.

In our context, it would be very useful to obtain a better understanding of the visual features found to be correlated to gene expression, since these automatically inferred visual features make up the phenotype of interest. As opposed to features detected by human experts which are much more subjective and much harder to quantify, these automatically derived features have a precise mathematical definition that allows a precise quantification. Their visualization would also enable a comparison with expert domain knowledge.

In contrast to fully connected network models, convolutional networks tend to be easier to interpret and visualize. We have found two types of visualizations to be particularly useful in our application.

1. The first looks for elements (pixels) of the original image that affect a given feature most. These can be determined using backpropagation of the feature of interest w.r.t. a given input image. In particular, we employ *guided backpropagation* [Springenberg, 2014], which tends to produce better visualizations by zeroing negative gradients during the standard backpropagation process through ReLU units. Such visualizations, which we denote as 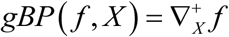 depend on the given input image *X* and were constructed for images from the *test dataset*, which is completely independent from the datasets used for model construction and respectively selection (training and respectively validation datasets).
2. The second visualization is independent of a specific input and amounts to determining a *synthetic input image* that maximizes the given feature 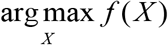 [Olah, 2017].

Note that unfortunately, guided backpropagation visualization cannot be applied exhaustively because it would involve an unmanageably large number of feature-histological image (*f,X*) pairs. To deal with this problem, we developed an algorithm for selecting a small number of representative histological image tiles to be visualized with guided backpropagation (see Supporting Information S1 File). The problem is also addressed by using the second visualization method mentioned above, which generates a single synthetic image per feature.

Guided backpropagation visualizations of select features are compared with synthetic images and immunohistochemistry stains for the genes found correlated with the selected feature.

Since the network features were optimized to aid in tissue discrimination, while many genes are also tissue-specific, it may be possible that certain gene-feature correlations are indirect via the tissue variable. To assess the prevalence of such indirect gene(*g*)-tissue(*t*)-feature(*f*) correlations, we performed conditional independence tests using partial correlations of the form *r*(*g,f* | *t*) with a significance threshold *p*=0.01.

## Results

First, we developed a classifier of histological images that is able to discriminate between 39 tissues of interest. A convolutional neural network enables learning visual representations that can be used as visual features in the subsequent stages of our analysis.

The resulting visual features are then quantified on an independent test dataset and correlated with paired gene expression data from the same subjects.

Finally, the features that are highly correlated to genes are visualized for better interpretability. The following sections describe the results of these analysis steps in more detail.

### Deep learning tissue classifiers achieve high accuracies

We trained and validated several different convolutional neural network architectures (from Table 1) on the GTEx image tile dataset. Table 2 shows the accuracies of the classifiers on both *validation* and *test* datasets, after training for 90 epochs on the *training* dataset. Note that the test dataset is completely independent, while the validation dataset has been used for model selection.

**Table 2.** Classification accuracies for image *tiles*.

| Network | accuracy (tiles) |  |
| --- | --- | --- |
|  | *validation* | *test* |
| AlexNet | 77.02% | 74.76% |
| VGG11 | 80.18% | 77.49% |
| VGG13 | 80.77% | 78.19% |
| VGG16 | 80.35% | 78.00% |
| VGG19 | 79.61% | 76.93% |
| VGG11_bn | 83.24% | 80.74% |
| VGG13_bn | 83.94% | 81.56% |
| VGG16_bn | 84.24% | 81.46% |
| VGG19_bn | 84.44% | 81.54% |
| ResNet34 | 82.51% | 80.09% |
| Inception_v3 | 81.13% | 78.99% |
| VGG16_1FC | 82.66% | 79.82% |
| VGG16_avg1FC | 83.79% | 80.92% |

As expected, AlexNet turned out to be the worst classifier among the ones tested. The various VGG architectures produced comparable results, regardless of their size. This may be due to the smaller number of classes (39) compared to ImageNet (1,000). On the other hand, batch normalization leads to improved classification accuracies, but also to representations that show much poorer correlation with gene expression data (see next section).

The modified architectures VGG16_1FC and VGG16_avg1FC perform slightly better than the original VGG16.

Overall, accuracies of 80-82% in identifying the correct tissue based on a single image tile are significant, but still not comparable to human performance. This is because single image tiles allow only a very limited field of view of the tissue.

However, as the images contain multiple tiles, we constructed a *tissue classifier for whole slide images (WSI)* by applying the tile classifier on all tiles of the given WSI and aggregating the corresponding predictions using the majority vote. Table 3 show the accuracies of the whole slide image classifiers obtained using the convolutional neural network models from Table 1.

**Table 3.** Classification accuracies for *whole slides*.

| Network | accuracy (slides) |  |
| --- | --- | --- |
|  | *validation* | *test* |
| AlexNet | 91.82% | 90.42% |
| VGG11 | 93.33% | 92.22% |
| VGG13 | 93.94% | 92.81% |
| VGG16 | 92.73% | 93.71% |
| VGG19 | 93.33% | 93.11% |
| VGG11_bn | 93.94% | 92.81% |
| VGG13_bn | 93.03% | 92.51% |
| VGG16_bn | 93.64% | 92.22% |
| VGG19_bn | 93.94% | 92.81% |
| ResNet34 | 93.33% | 92.51% |
| Inception_v3 | 92.12% | 92.22% |
| VGG16_1FC | 93.33% | 93.11% |
| VGG16_avg1FC | 93.64% | 94.01% |

### Histological features inferred by VGG architectures correlate well with gene expression data

We calculated the correlations between the visual features of convolutional networks trained on histopathological images and paired gene expression data. We found numerous genes whose expression is correlated with many visual histological features.

Table 4 shows the numbers of significant gene-feature pairs (as well as unique genes, features and respectively tissues involved in these relations) for the various CNN architectures using fixed correlation-(*R*_*T*_ = 0.8) and *log*_*2*_ expression thresholds *E*_*T*_ = 10 (where *log*_*2*_(1+*E*) ≤ *E*_*T*_). Histograms of features, genes and their correlations are shown in S1 Fig.

**Table 4.** Numbers of significantly correlated gene-feature pairs for various network architectures. Fixed correlation- and *log*_*2*_ gene expression thresholds are used: *R*_*T*_ = 0.8, *E*_*T*_ = 10. Numbers of unique genes, features and tissues involved are also shown (test dataset).

| Network | gene-feature pairs | unique genes | unique features | unique tissues |
| --- | --- | --- | --- | --- |
| AlexNet | 424 | 55 | 112 | 11 |
| VGG11 | 1,175 | 70 | 204 | 12 |
| VGG13 | 2,149 | 70 | 307 | 13 |
| VGG16 | 2,176 | 74 | 346 | 13 |
| VGG19 | 1,227 | 71 | 321 | 14 |
| VGG11_bn | 34 | 19 | 7 | 3 |
| VGG13_bn | 59 | 22 | 17 | 6 |
| VGG16_bn | 51 | 20 | 11 | 4 |
| ResNet34 | 3 | 2 | 3 | 2 |
| Inception_v3 | 23 | 12 | 7 | 2 |
| VGG16_1FC | 2,714 | 69 | 363 | 13 |
| VGG16_avg1FC | 3,114 | 83 | 448 | 15 |

Note that although batch normalization slightly improved *tile* classification accuracies (but not *WSI* classification accuracies), it also produced visual representations (features) with dramatically poorer correlation with gene expression data. For example, for VGG16\_bn, we could identify only 51 significant gene-feature pairs, involving just 20 unique genes and 11 unique features, compared to 2,176 gene-feature pairs implicating 74 unique genes and 346 unique features for VGG16 without batch normalization.

Unless stated otherwise, we concentrate in the following on visual representations obtained using the VGG16 architecture. (The results for AlexNet and the other VGG architectures without batch normalization are similar.)

VGG16 was selected not just based on it achieving one of the highest WSI classification accuracies on the test dataset (93.71%, Table 3), but also since its last convolutional layer is not directly connected to the output classes (as in the case of VGG16_1FC and VGG16_avg1FC). This is assumed to enable a higher degree of independence of the final convolutional layers from the output classes, rendering them more data oriented.

Table 5 shows the numbers of significantly correlated gene-feature pairs for various correlation-and *log*_*2*_ gene expression thresholds (for the VGG16 network).

**Table 5.** Numbers of significantly correlated gene-feature pairs for various correlation- and *log*_*2*_ gene expression thresholds. Numbers of unique genes, features and tissues involved are also shown (VGG16 network, test dataset).

| Correlation threshold | $\log_2$ expression threshold | gene-feature pairs | unique genes | unique features | unique tissues |
| --- | --- | --- | --- | --- | --- |
| 0.8 | 10 | 2,176 | 74 | 346 | 13 |
| 0.75 | 10 | 4,308 | 115 | 647 | 22 |
| 0.7 | 10 | 7,984 | 213 | 1,146 | 28 |
| 0.75 | 8 | 9,624 | 312 | 926 | 26 |
| 0.8 | 7 | 8,055 | 365 | 576 | 18 |
| 0.75 | 7 | 15,062 | 535 | 1,046 | 31 |
| 0.7 | 7 | 27,671 | 995 | 1,947 | 36 |

Using a correlation threshold *R*_*T*_ = 0.7 and a *log*_*2*_ expression threshold *E*_*T*_ = 7, we obtained 27,671 significant correlated gene-feature pairs involving 995 unique genes and 1,947 features (for the VGG16 architecture and the test dataset). We call these gene-feature pairs significant, because the p-values computed using permutation tests (with *N*=1,000 permutations) were all *p*<10^−3^.

Fig 3 shows the numbers of significantly correlated genes for the 31 layers of VGG16 (numbered 0 to 30). All features (channels) of a given layer were aggregated, since VGG16 has a too large number of features (9,920 in total). (See also S2 Fig (A) for numbers of correlated genes for *individual* features.) The correlated genes were broken down according to their tissues of maximal expression. Note that the highest numbers of correlated genes correspond to layers with non-negative output (ReLU or MaxPool2d – cf. layer numbers in the VGG16 entry in Table 1).

**Fig 3.**
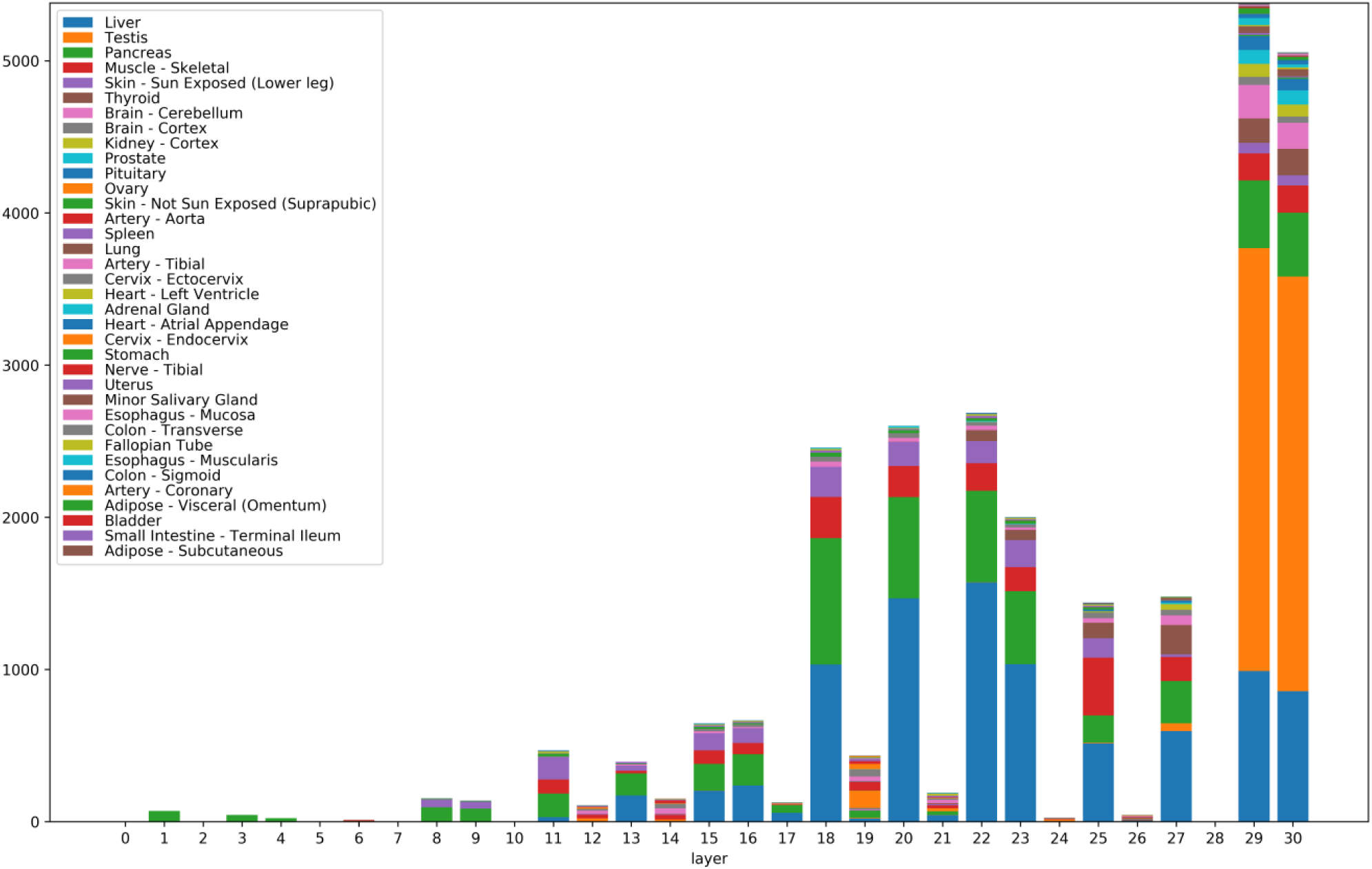
Numbers of significantly correlated genes for the 31 layers of VGG16. Significantly correlated genes were aggregated for all features (channels) belonging to a given layer (i.e. for all features of the form [layer]_[channel]). The correlated genes were broken down according to their tissues of maximal expression. Tissues are color-coded. Colors of the figure bars scanned bottom-up correspond to colors in the legend read top to bottom.

We also remark that the features with significant gene correlations do not necessarily belong to the final layers of the convolutional neural network, even though one may have expected the final layers to be better correlated with the training classes (i.e. the tissue types). Also note that certain tissues with a simpler morphology, such as liver (blue in Fig 3) show correlated genes with features across many layers of the network, from lower levels to the highest levels. Other morphologically more complex tissues, such as testis (orange in Fig 3) express genes correlated almost exclusively with the final layers 29 and 30 (which are able to detect such complex histological patterns). See also examples of histological images for these tissues in Fig 6 (column 1, rows 1 and 2).

For an increased *specificity* and in order to reduce the number of gene-feature pairs subject to human evaluation, we initially performed an analysis with higher thresholds *R*_*T*_ = 0.8 and *E*_*T*_ = 10 (resulting in 2,176 gene-feature pairs, involving 74 genes and 346 features).

A second analysis with lower thresholds (*R*_*T*_ = 0.75, *E*_*T*_ = 7) was performed with the aim of improving the *sensitivity* of the initial analysis (15,062 gene-feature pairs, involving 535 genes and 1,046 features -S2 Table).

Since gene-feature *correlations* do not necessarily indicate *causation* (i.e. the genes involved are not necessarily causal factors determining the observed visual feature), we tried to select a subset of genes, based on known annotations that are more likely to play a causal role in shaping the histological morphology of the tissues. In the following, we report on the subset of genes with Gene Ontology ‘*developmental process*’ or ‘*transcription regulator activity*’ annotations (either direct, or inherited) [Ashburner, 2000; Gene Ontology Consortium, 2019] (Table 6 and S3 Table). While some developmental genes are turned off after completion of the developmental program, many continue to be expressed in adult tissue, to coordinate and maintain its proper structure and function.

**Table 6.**
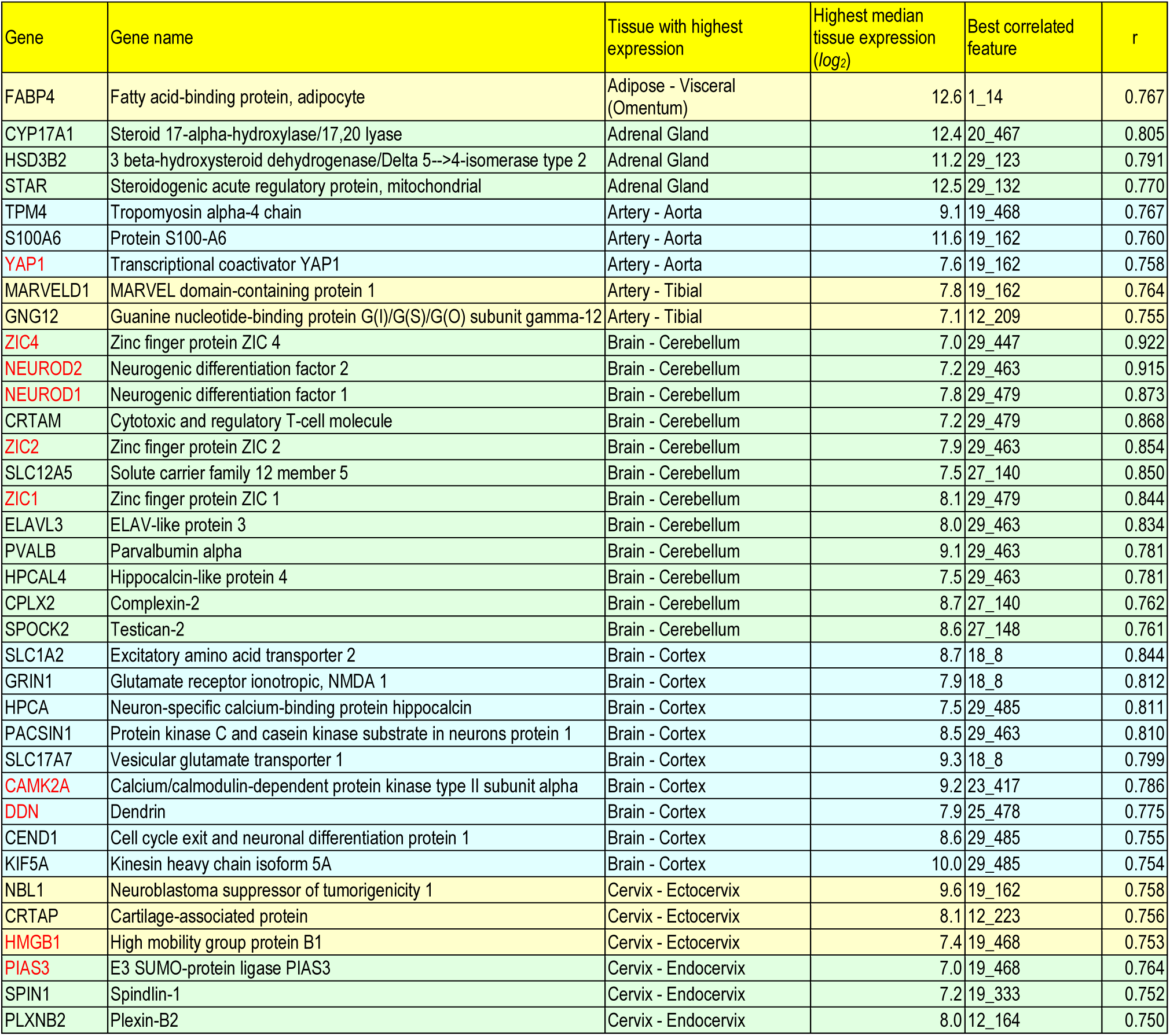
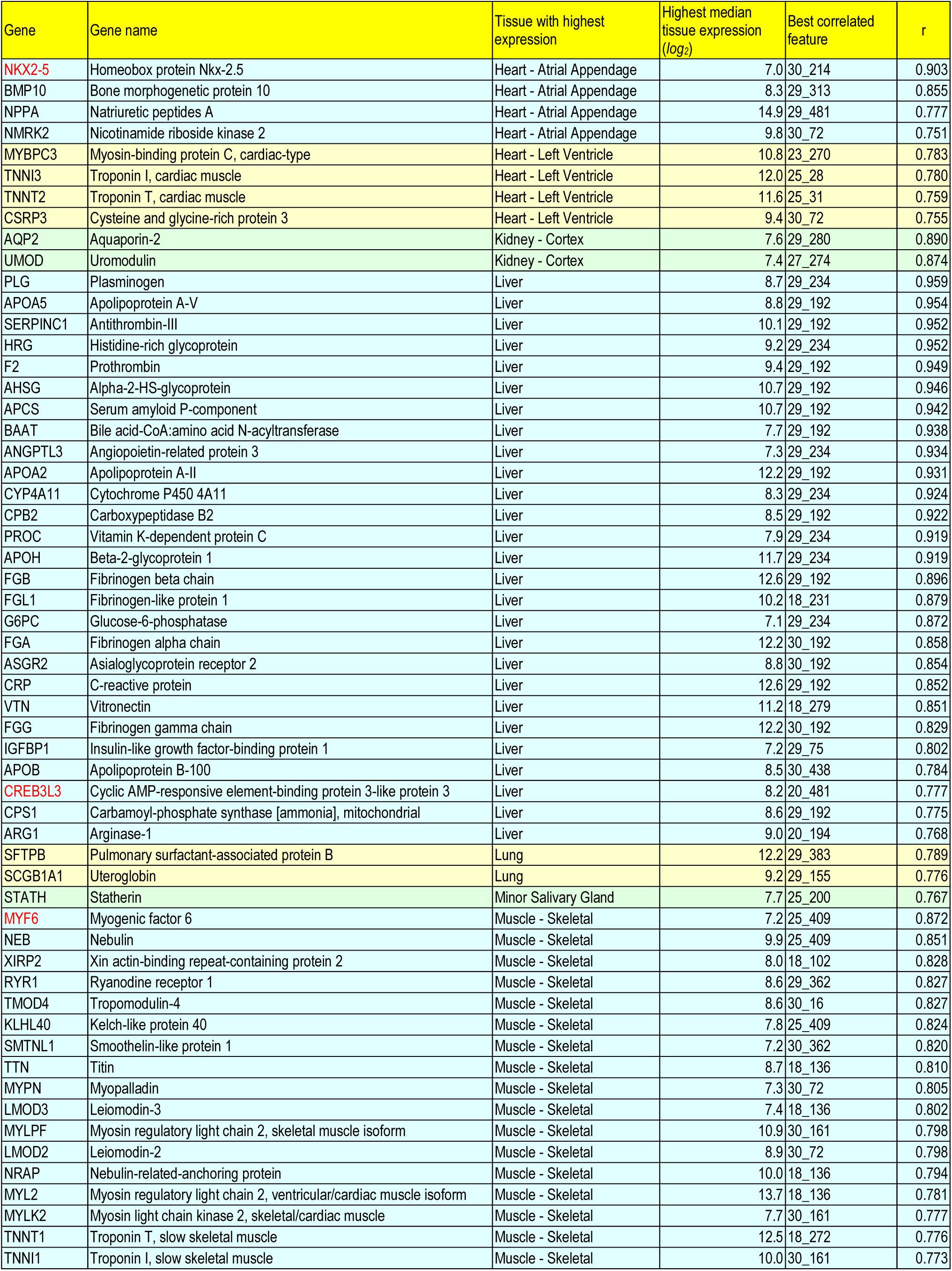
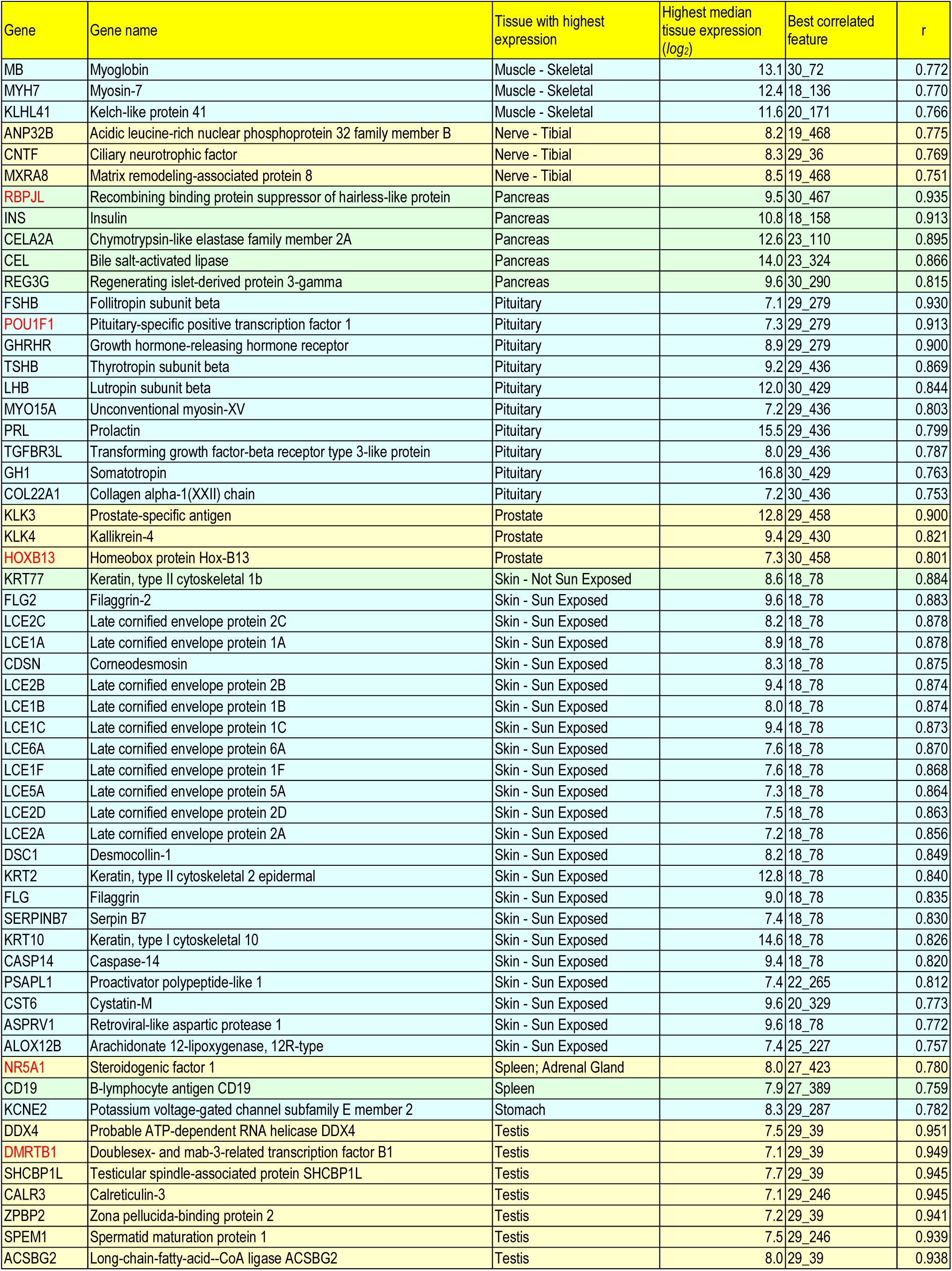
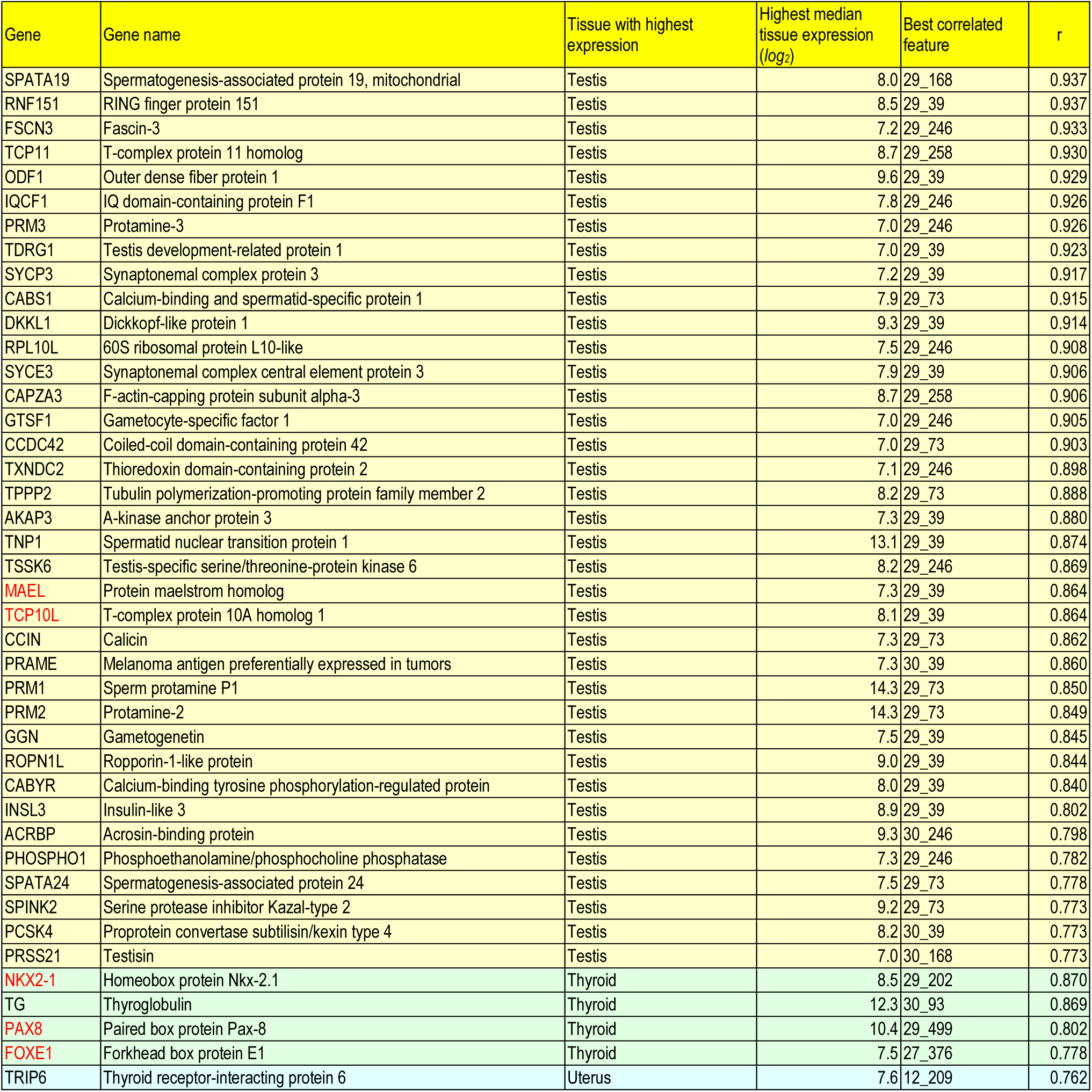
Developmental and transcription regulation genes correlated with visual features. Genes with Gene Ontology ‘*developmental process*’ or ‘*transcription regulator activity*’ annotations are grouped w.r.t. tissues and ordered by correlation - for each gene we only show the best correlated feature, in the form [layer]_[channel] (for the architecture VGG16 and the test dataset; correlation threshold=0.75, *log*_*2*_ gene expression threshold=7). Genes with ‘*transcription regulator activity*’ are shown in red.

Many of the genes significantly correlated with histological features have crucial roles in the development and maintenance of the corresponding tissues. The transcription factors ZIC1, ZIC2, ZIC4, NEUROD1 and NEUROD2 are known to be involved in brain and more specifically cerebellar development [Aruga, 2018; Pieper, 2019], NKX2-5 and BMP10 are implicated in heart development [Akazawa, 2005; Chen, 2004], POU1F1 regulates expression of several genes involved in pituitary development and hormone expression [Kelberman, 2009], NKX2-1, PAX8 and FOXE1 are key thyroid transcription factors, with a fundamental role in the proper formation of the thyroid gland and in maintaining its functional differentiated state in the adult organism [Fernandez, 2015], etc. (Table 6, S3 Table). While a correlational approach such as the present one cannot replace perturbational tests of causality involved in shaping tissue morphology, it can provide appropriate candidates for such tests.

We also studied the reproducibility of gene-feature correlations w.r.t. the dataset used. More precisely, we determined the Pearson correlation coefficient between the (Fisher transformed) gene-feature correlations computed w.r.t. the validation and respectively test datasets (S3 Fig):

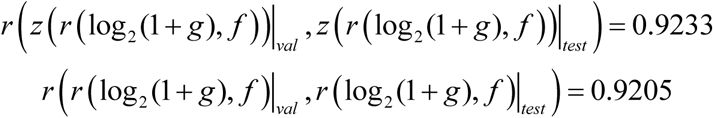

where *z*(*r*) is the Fisher transform of correlation *r*. Gene-feature correlations thus show a good reproducibility across datasets.

### Visualization of histological features

To investigate the biological relevance of the discovered gene-feature correlations, we analyzed in detail different methods of visualizing the features inferred by convolutional networks. We applied two different visualization methods. The first based on ‘*guided backpropagation*’ determines the regions of a histopathological image that affect the feature of interest most. However, this visualization method cannot be applied exhaustively because it would involve an unmanageably large number of feature-histological image pairs. To deal with this problem, we developed an algorithm for selecting a small number of representative histological image tiles to be visualized with guided backpropagation (S1 File). We also considered a second feature visualization method that is independent of any input image. The method involves generating a synthetic input image that optimizes the network response to the visual feature of interest. Such synthetic images look similar to real histological images that strongly activate the visual feature of interest.

Figures 4-6 show such visualizations of select histological features. Note that *guided backpropagation* (column 2) emphasizes important structural features of the original histological images (column 1). For example, performing guided backpropagation (gBP) of feature 29_499 on a thyroid sample produces patterns that clearly overlap known histological structures of thyroid tissues, namely follicular cells in blue and follicle boundaries in yellow (*row 3*, column 2 of Fig 5). More precisely, the blue “dots” in the gBP image precisely overlap the follicular cells, while the yellow patterns correspond to boundaries of follicles (please compare with the original image from column 1).

**Fig 4.**
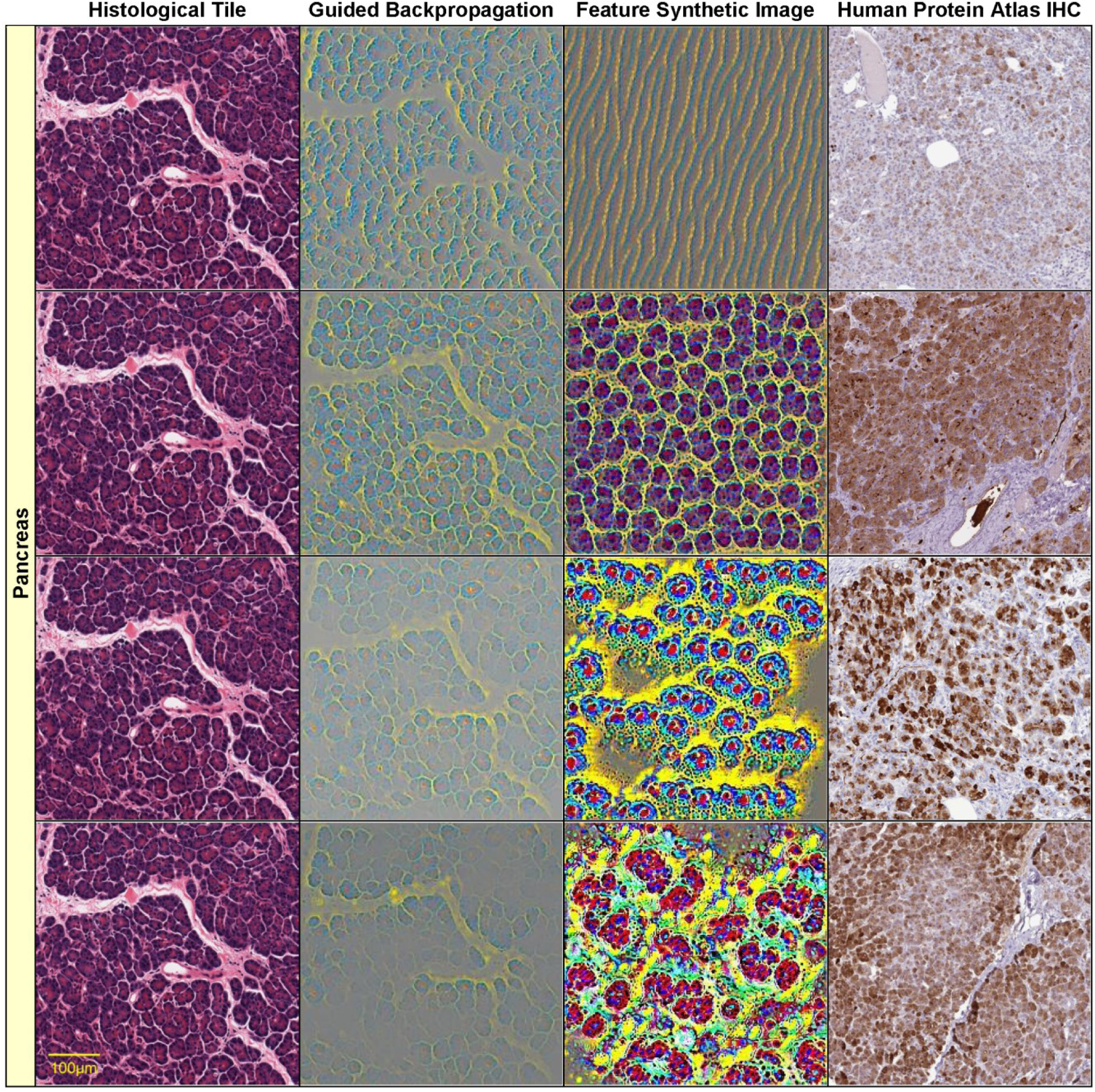
Visualizations of select histological features. The following features (of the form [layer]_[channel]) found correlated with specific genes are visualized on each row: row 1: 8_55 - CTRC (*r* = 0.85) row 2: 18_13 - CUZD1 (*r* = 0.866) row 3: 23_324 - CELA3B (*r* = 0.868) row 4: 30_467 - AMY2A (*r* = 0.928) Original image (column 1), guided backpropagation of the feature on the original image (column 2), synthetic image of the feature (column 3), immunohistochemistry image for the corresponding gene from the Human Protein Atlas (column 4). All visualizations are for *pancreas* sample tile GTEX-11NSD-0526_32_5.

**Fig 5.**
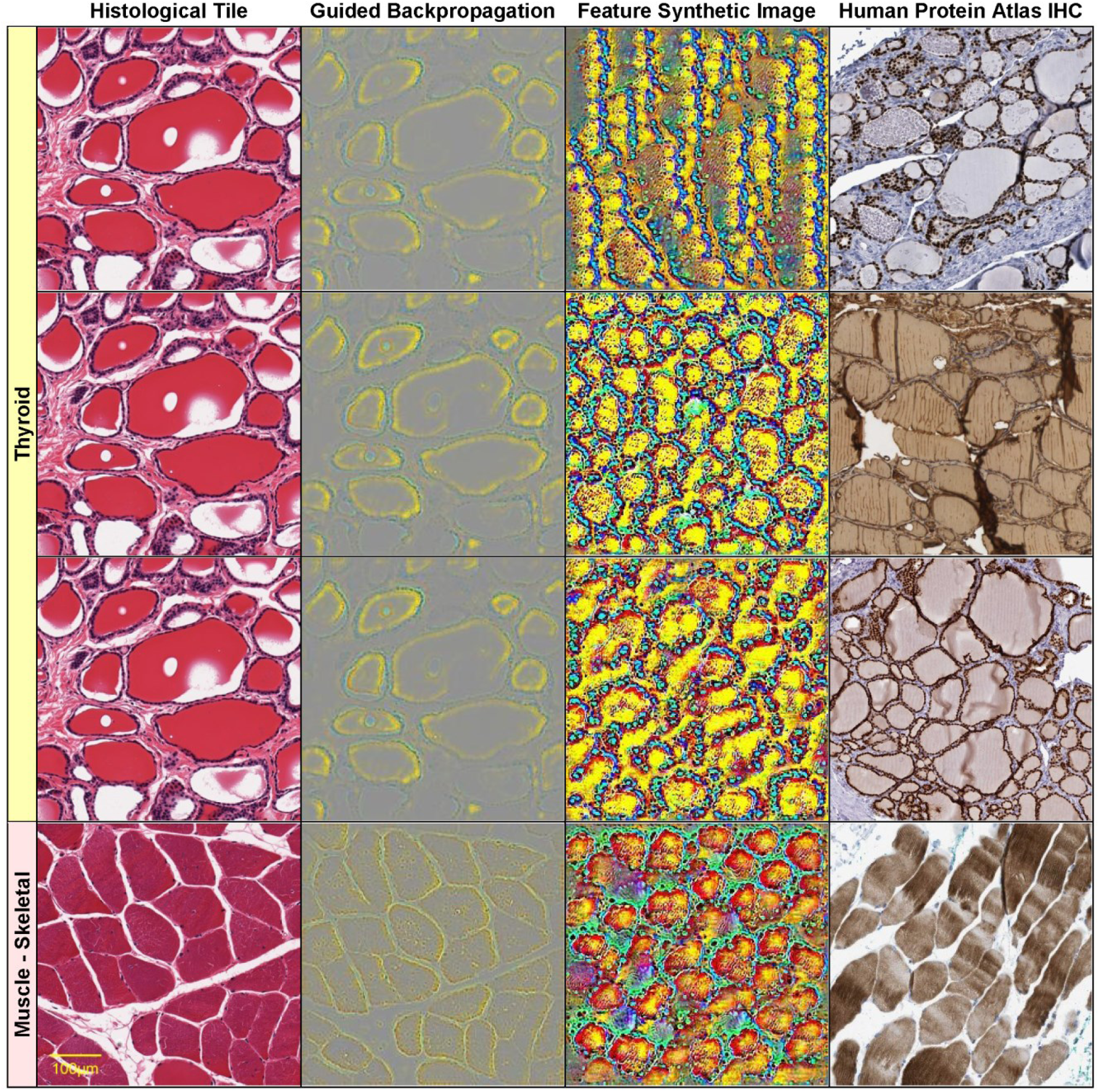
Visualizations of select histological features. The following features (of the form [layer]_[channel]) found correlated with specific genes are visualized on each row: row 1: 22_405 NKX2-1 (*r* = 0.726) Thyroid GTEX-11NSD-0126_31_16 row 2: 27_343 TG (*r* = 0.801) Thyroid GTEX-11NSD-0126_31_16 row 3: 29_499 PAX8 (*r* = 0.802) Thyroid GTEX-11NSD-0126_31_16 row 4: 25_260 NEB (*r* = 0.831) Muscle - Skeletal GTEX-145ME-2026_39_19 Original image (column 1), guided backpropagation of the feature on the original image (column 2), synthetic image of the feature (column 3), immunohistochemistry image for the corresponding gene from the Human Protein Atlas (column 4).

**Fig 6.**
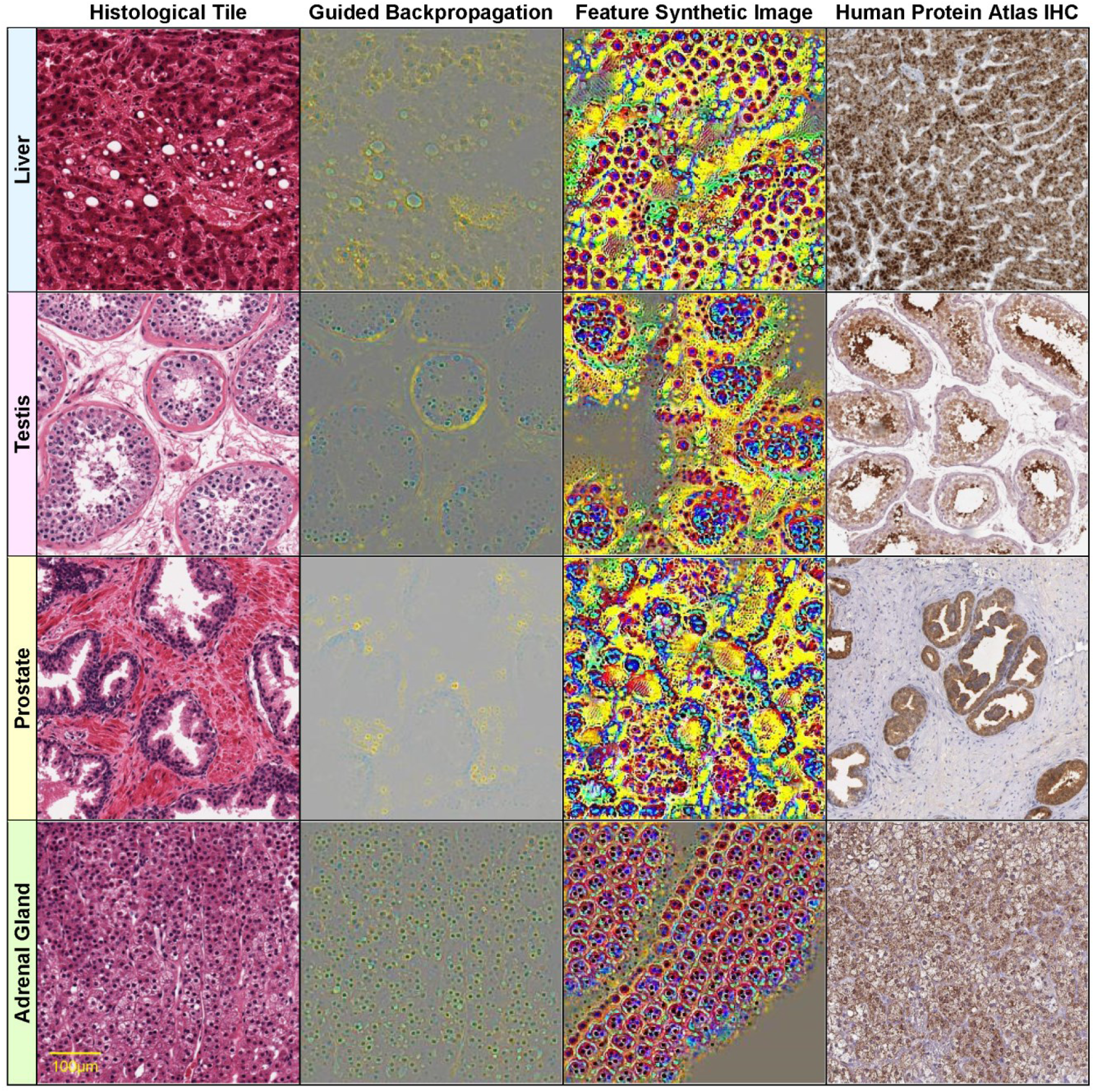
Visualizations of select histological features. The following features (of the form [layer]_[channel]) found correlated with specific genes are visualized on each row: row 1: 29_234 AGXT (*r* = 0.939) Liver GTEX-Q2AG-1126_9_7 row 2: 29_118 CALR3 (*r* = 0.829) Testis GTEX-11NSD-1026_5_27 row 3: 30_244 KLK3 (*r* = 0.828) Prostate GTEX-V955-1826_8_13 row 4: 20_137 CYP11B1 (*r* = 0.850) Adrenal Gland GTEX-QLQW-0226_25_9 Original image (column 1), guided backpropagation of the feature on the original image (column 2), synthetic image of the feature (column 3), immunohistochemistry image for the corresponding gene from the Human Protein Atlas (column 4).

On the other hand, column 3 displays a synthetically generated image that maximizes the corresponding feature (29_499). The image was generated by optimization of the feature using incremental changes to an initially random input image. Still, the synthetic image shows a remarkable resemblance to the histological structure of thyroid tissue (except for color differences due to color normalization).

We also show in column 4 the expression of PAX8 in an immunohistochemistry (IHC) stain of thyroid tissue from the Human Protein Atlas [Uhlén, 2015]. Note that PAX8 was identified as a correlate of the above-mentioned feature (29_499) with correlation *r*=0.802 (Table 6) and is specifically expressed in the follicular cells (corresponding to the blue “dots” in the gBP image from column 3). PAX8 encodes a member of the paired box family of transcription factors and is known to be involved in thyroid follicular cell development and expression of thyroid-specific genes. Although PAX8 is expressed during embryonic development and is subsequently turned off in most adult tissues, its expression is maintained in the thyroid [Fernandez, 2015].

Note that the features with significant gene correlations do not necessarily belong to the final layers of the convolutional neural network (as one may expect the final layers to be better correlated with the training classes, i.e. the tissue types). Fig 4 illustrates this by showing four features *at various layers* (8, 18, 23 and 30) of the VGG16 architecture (which has in total 31 layers, numbered 0 to 30) for a pancreas slide. Lower layers tend to capture simpler visual features, such as blue/yellow edges in the synthetic image of feature 8_55 (row 1, column 3 of Fig 4). In the corresponding guided backpropagation image (column 2), these edges are sufficient to emphasize the yellow boundaries between the blue pancreatic acinar cells. Higher layer features detect more complex histological features, such as round “cell-like” structures that tile the entire image (feature 18_13, row 2, column 3 of Fig 4), or more complex blue cell-like structures (including red nuclei) separated by yellow interstitia (feature 23_324, row 3, column 3 of Fig 4). The highest level feature, 30_467 (row 4, column 3 of Fig 4) seems to detect even more complex patterns involving mostly yellow connective tissue and surrounding red/blue “cells”.

Other examples of visualizations of features from different layers are shown in rows 1-3 of Fig 5 for a thyroid slide. The lower level feature 22_405 detects boundaries between blue follicular cells and yellow follicle lumens, while the higher level features 27_343 and 29_499 are activated by more complex, round shapes of blue follicular cells surrounding yellow-red follicle interiors. All of these features are significantly correlated with thyroglobulin TG expression *r*(TG,22_405)=0.805, *r*(TG,27_343)=0.801, *r*(TG,29_499)=0.781, but also with other key thyroidal genes, such as TSHR, NKX2-1, PAX8 and FOXE1: *r*(TSHR,22_405)=0.828, *r*(TSHR,27_343)=0.819, *r*(TSHR,29_499)=0.824, *r*(FOXE1,22_405)=0.744, *r*(NKX2-1,22_405)= 0.726, *r*(NKX2-1,29_499)=0.711, *r*(PAX8,29_499)=0.802. Note that NKX2-1, FOXE1 and PAX8 are three of the four key thyroid transcription factors, with a fundamental role in the proper formation of the thyroid gland and in maintaining its functional differentiated state in the adult organism [Fernandez, 2015]. The fourth thyroid transcription factor, HHEX, is missing from our analysis due to its expression slightly below the *log*_*2*_ expression threshold used (the median HHEX expression is 6.84, just below the threshold 7). For illustration purposes, we show, in column 4 of Fig 5, IHC stains for distinct thyroid genes (NKX2-1 for feature 22_405, TG for 27_343 and respectively PAX8 for 29_499). Note that all of these IHC images are remarkably similar to the corresponding guided backpropagation (gBP) images from column 2 and the synthetic images from column 3.

It is remarkable that spatial expression patterns of genes, as assessed by IHC, are frequently very similar to the gBP images of their correlated features. Still, not all significantly correlated gene-feature pairs display such a good similarity between the gBP image of the feature and the spatial expression pattern of the gene. This is due to the complete lack of spatial specificity of the RNA-seq gene expression data, as well as to the rather coarse-grained classes used for training the visual classifier – just tissue labels, without spatial annotations of tissue substructure. This is the case of tissue-specific genes, which may display *indirect gene-tissue-feature correlations* with distinct spatial specificities of the gene-tissue and respectively feature-tissue correlations. (The gene may be specifically expressed in a certain tissue substructure, while the visual feature may correspond to a distinct substructure of the same tissue. As long as the two different tissue substructures have similar distributions in the tissue slides, we may have indirect gene-tissue-feature correlations without perfect spatial overlap of gene expression and the visual feature.)

For example, the high-level feature 30_467 seems to detect primarily yellow connective tissue in pancreas samples, rather than acinar cells, which express the AMY2A gene (row 4, column 2 of Fig 4). The partial correlation *r*(*g,f* | *t*) = 0.107 is much lower than *r*(*g,f*) = 0.928, with a p-value of the conditional independence test *p*=0.0501 > 0.01. Therefore, the AMY2A expression and the feature 30_467 are independent conditionally on the tissue (i.e. their correlation is indirect via the tissue variable).

To assess the prevalence of such indirect gene(*g*)-tissue(*t*)-feature(*f*) correlations, we performed conditional independence tests using partial correlations of the form *r*(*g,f* | *t*) with a significance threshold *p*=0.01. Table 7 shows the resulting numbers of indirect dependencies for two different significance thresholds of the conditional independence test (*α*=0.01 and 0.05). The gene-tissue-feature (*g*-*t*-*f*) case corresponds to the indirect gene-feature correlations mediated by the tissue variable, discussed above. Indirect feature-gene-tissue (*f*-*g*-*t*) dependencies correspond to genes that mediate the feature-tissue correlation (for which the classifier is responsible). There are significantly fewer such indirect dependencies, and even fewer ones of the form *g*-*f*-*t*.

**Table 7.**
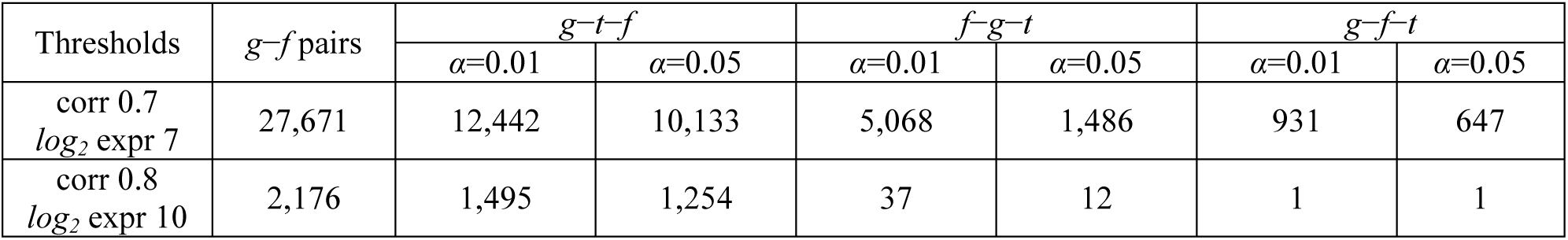
Numbers of significant gene-feature pairs with indirect dependencies gene-tissue-feature (*g*-*t*-*f*), feature-gene-tissue (*f*-*g*-*t*), gene-feature-tissue (*g*-*f*-*t*). Conditional independence tests with α=0.01 and respectively α=0.05.

| Thresholds | $g$-$f$ pairs | $g$-$t$-$f$ | | $f$-$g$-$t$ | | $g$-$f$-$t$ | |
| --- | --- | --- | --- | --- | --- | --- | --- |
| | | $\alpha = 0.01$ | $\alpha = 0.05$ | $\alpha = 0.01$ | $\alpha = 0.05$ | $\alpha = 0.01$ | $\alpha = 0.05$ |
| corr 0.7<br>$\log_2$ expr 7 | 27,671 | 12,442 | 10,133 | 5,068 | 1,486 | 931 | 647 |
| corr 0.8<br>$\log_2$ expr 10 | 2,176 | 1,495 | 1,254 | 37 | 12 | 1 | 1 |

The visualizations of the prostate sample from row 3 of Fig 6 illustrate an interesting intermediary case in which although conditioning on the tissue leads to a big drop in correlation (from *r*(*g,f*)=0.828 to *r*(*g,f* | *t*)=0.167), the conditional independence test still rejects the null hypothesis (*p*=0.00215 < 0.01, i.e. there is still conditional *dependence*, although a weak one). The gene-feature correlation is therefore not entirely explainable by the indirect gene-tissue-feature influence. This can also be seen visually in the gBP image of feature 30_244, which detects blue glandular cells and surrounding yellow stroma (row 3, column 2 of Fig 6), similar to the spatial expression pattern of the KLK3 gene, which is specifically expressed in the glandular cells (see IHC image from column 4). The imperfect nature of the dependence is due to the feature detecting mostly basally situated glandular cells (blue) rather than all glandular cells.

## Discussion

Due to the limited numbers of high-quality datasets comprising both histopathological images and genomic data for the same subjects, there are very few research publications trying to combine visual features with genomics. Also, the relatively small numbers of samples compared to genes and image features make such integration problems difficult (‘small sample problem’).

The sparse canonical correlation analysis (CCA) used in [Ash, 2018] is an elegant and very general technique of determining correlations between two *sets* of variables (more precisely, finding linear combinations of variables from each of the two sets that are maximally correlated to one another). Since the sets of genes and respectively image features are very large, CCA is ideal for determining *general trends* (CCA components) in the complicated correlation structure of genes and features. On the other hand, in this paper we are interested in determining *individual* visual features correlated with genes, rather than CCA *composites* of such features, which may be hard to visualize and interpret. Such individual visual features might also be essential for discriminating between visually similar tissues (especially in the case of the number of variables exceeding the number of samples – sparsity constraints mitigate this problem, but do not eliminate it altogether). Moreover, multivariate regression analysis is typically not recommended for small samples. Also, while we use spatially invariant feature encodings obtained by aggregating feature values over both *spatial* and *tile* dimensions, [Ash, 2018] average their feature encodings only over the tile (“window”) dimension and not over the spatial dimensions.

Other work searched for relationships between histological images and somatic cancer mutations. For example, [Coudray, 2018] trained a CNN to predict the mutational status of just 10 genes (the most frequently mutated genes in lung cancer, rather than a genome-wide study). Similar mutation predictors were developed for hepatocellular carcinoma [Chen, 2020] and prostate cancer [Schaumberg, 2018]. But the focus of these studies was obtaining mutation predictors, rather than correlating the mutational status with visual histological features.

There is a much larger body of research on purely visual algorithms for analyzing histological slides (without considering correlations with transcriptomics). However, a direct comparison of tissue classification accuracies is probably not very meaningful, since different sets of tissues were used in most papers.

The GTEx dataset was also used to obtain tissue classifiers in [Bizzego, 2019], for 5, 10, 20 and respectively 30 tissues. The highest *tile classification accuracies* reported in [Bizzego, 2019] for 30 tissues are 61.8% for VGG pretrained on Imagenet and respectively 77.1% for the network retrained from scratch on histological images, compared to our 81.5% accuracy for classifying 39 tissues (30 tissue *WSI* classification accuracies are not reported in [Bizzego, 2019]). The dataset in [Bizzego, 2019] contains 53,000 tiles obtained from 787 WSI images from GTEx, for 30 tissues. However, as already mentioned above, some of the additional tissues in our extended set of 39 tissues are hard to distinguish (e.g. ‘Artery – Aorta’ and ‘Artery – Coronary’ versus ‘Artery – Tibial’).

While many researches have already covered normal and diseased tissue classification in digital histopathology, only a small minority have begun to address the problem of interpretability of the resulting models. From this perspective, there are two main contributions of this work, one dealing with the *visual interpretability* of the models, the second involving their *biological interpretability*.

Firstly, we develop *visualization methods* for the features that are inferred automatically by deep learning architectures. These histological features have certain domain-specific characteristics, namely they are *location independent*, as well as precisely *quantifiable*. Quantifiability involves being able to estimate the numbers of occurrences of a specific feature in a given histological image, such as the numbers of specific cellular structures in a slide.

The second main contribution consists in assessing the *biological interpretability* of the deep learning models by correlating their inferred features with matching gene expression data. Such features correlated with gene expression have more than a visual interpretation – they correspond to biological processes at the level of the genome.

It is remarkable that the synthetic images of certain features resemble the corresponding tissue structures extremely well. Genes predominantly expressed in tissues with simpler morphologies tend to be correlated with simpler, lower level visual features, while genes specific to more complex tissue morphologies correlate with higher level features.

However, not all CNN architectures infer features that tend to be well correlated with gene expression profiles, for example VGG networks with batch normalization, Inception_v3 or ResNet (which need batch normalization to deal with their extreme depth). It seems that batch normalization produces slight improvements in tile classification accuracies (though not for *whole* slides) at the expense of biological interpretability (i.e. significantly worse correlations of inferred features with the transcriptome – see Table 4).

## Conclusions

Current artificial intelligence systems are still not sufficiently developed to fully take over the tasks of the histopathologist, who is ultimately responsible for the final clinical decision. But the AI system could prove invaluable in assisting the clinical decision process. To do so, it needs to provide the histologist with as much meaningful knowledge as possible. Unfortunately, there is at present no established way to easily explain why a specific decision was made by a network when dealing with a given histopathology image. This is generally unacceptable in the medical community, as clinicians typically need to understand and justify the reasons for a specific decision. A reliable diagnosis must be transparent and fully comprehensible. This is also especially important for obtaining regulatory approval for use in clinical practice [Tizhoosh, 2018]. Of particular concern in the medical field is the uncertainty of the decisions taken by a deep network, which can be radically affected even by the change of very few pixels in an image, in case of a so-called adversarial attack [Athalye, 2018].

Visualizations of the features automatically inferred by a deep neural network are a first step toward making the decisions of the network more *interpretable* and *explainable*. Moreover, correlating the visual histological features with specific gene expression profiles increases the confidence in the *biological interpretability* of these features, since the genes and their expression are responsible for the cell structures that make up these visual features.

This paper deals with identifying transcriptomic correlates of histology using Deep Learning, at first just in normal tissues, as a small step towards bridging the wide explanatory gap between genes, their expression and the complex cellular structures of tissues that make up histological phenotypes. Of course, the relationships discovered in this study are only correlational. For elucidating causality, more complicated, perturbational experiments are needed, but the methodology used here is still applicable.

## Acknowledgments

This work was partially supported by the projects PN 1937-0601/2019 and PN 1937-0301/CPN 301 300/2019.

We are also deeply grateful to the GTEx consortium for making the gene expression data and histological images publicly available. The Genotype-Tissue Expression (GTEx) Project was supported by the Common Fund of the Office of the Director of the National Institutes of Health, and by NCI, NHGRI, NHLBI, NIDA, NIMH, and NINDS. The data used for the analyses described in this manuscript were obtained from the GTEx Portal (https://gtexportal.org/home/).

## Author Contributions

**Conceptualization:** Liviu Badea

**Data curation:** Liviu Badea, Emil Stanescu

**Formal analysis:** Liviu Badea

**Investigation:** Liviu Badea, Emil Stanescu

**Methodology:** Liviu Badea

**Project administration:** Liviu Badea

**Software:** Liviu Badea

**Supervision:** Liviu Badea

**Validation:** Liviu Badea, Emil Stanescu

**Visualization:** Liviu Badea, Emil Stanescu

**Writing – original draft:** Liviu Badea

**Writing – review & editing:** Emil Stanescu

## Abbreviations

H&E: hematoxylin and eosin
WSI: Whole Slide Image
GTEx: Genotype-Tissue Expression project
DL: Deep Learning
CNN: Convolutional Neural Network
gBP: guided Backpropagation
IHC: immunohistochemistry

## Supporting information

**S1 Fig.**
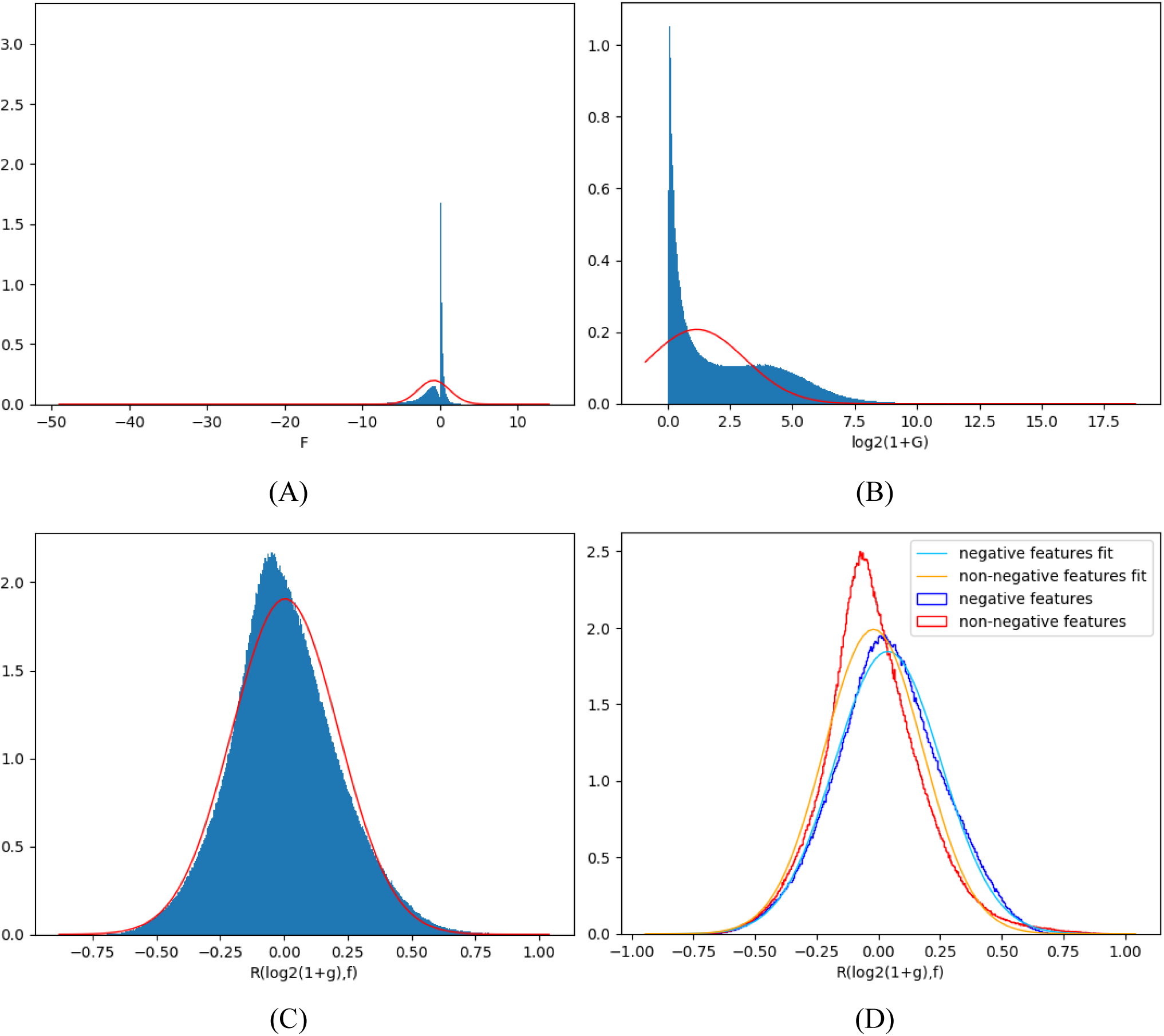
Histograms of features, genes and their correlation. (A) Histogram of feature values. (B) Histogram of *log*_*2*_ transformed gene expression values *log*_*2*_(1+*g*). (C) Histogram of gene-feature correlations (for genes with highest median tissue *log*_*2*_ expression over 10). (D) Histogram of gene-feature correlations separately for non-negative features (e.g. outputs of ReLU units) and potentially negative features.

**S2 Fig.**
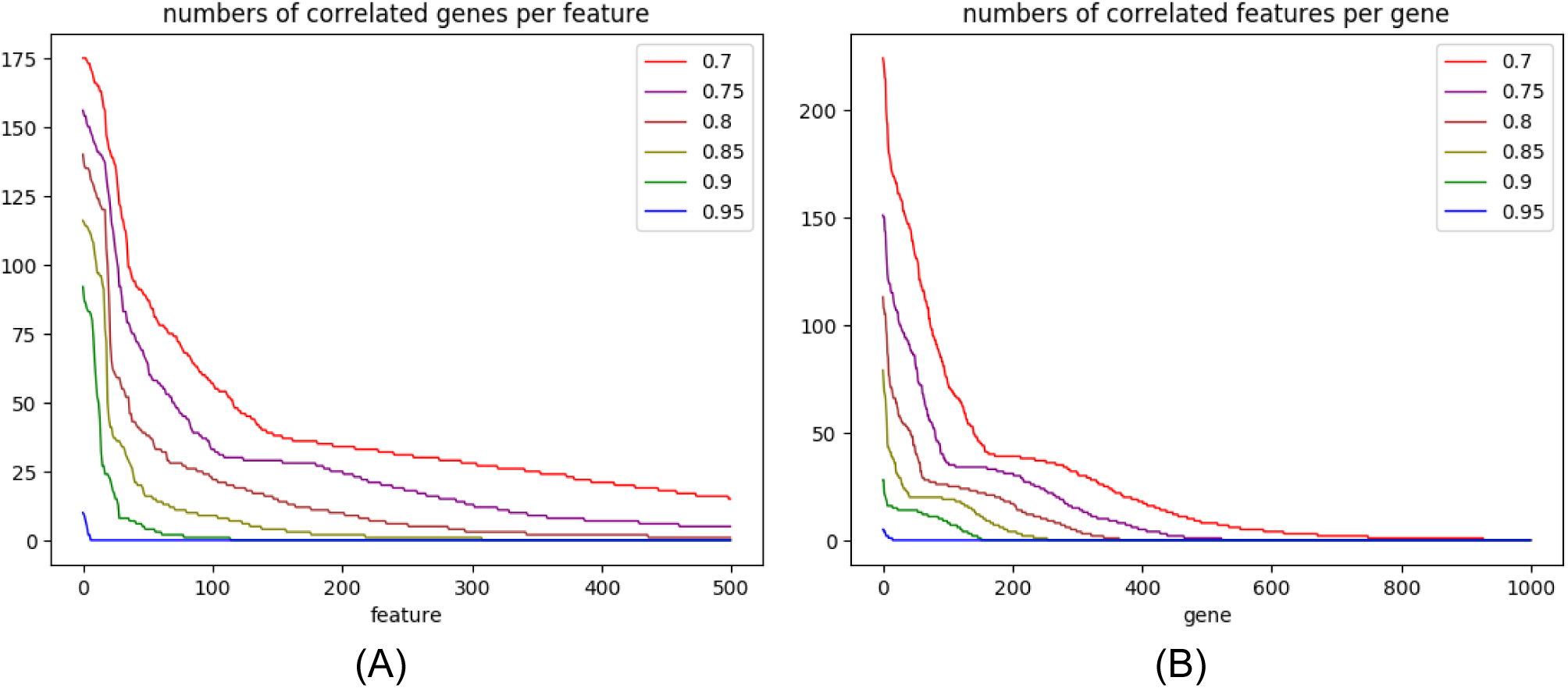
Numbers of correlated genes for individual features and respectively correlated features per gene. (A) Numbers of correlated genes for individual features. Features are sorted in decreasing order of the corresponding numbers of correlated genes. (B) Numbers of correlated features per gene. Genes are sorted in decreasing order of the corresponding numbers of correlated features. Various correlation thresholds are applied (from 0.7 to 0.95).

**S3 Fig.**
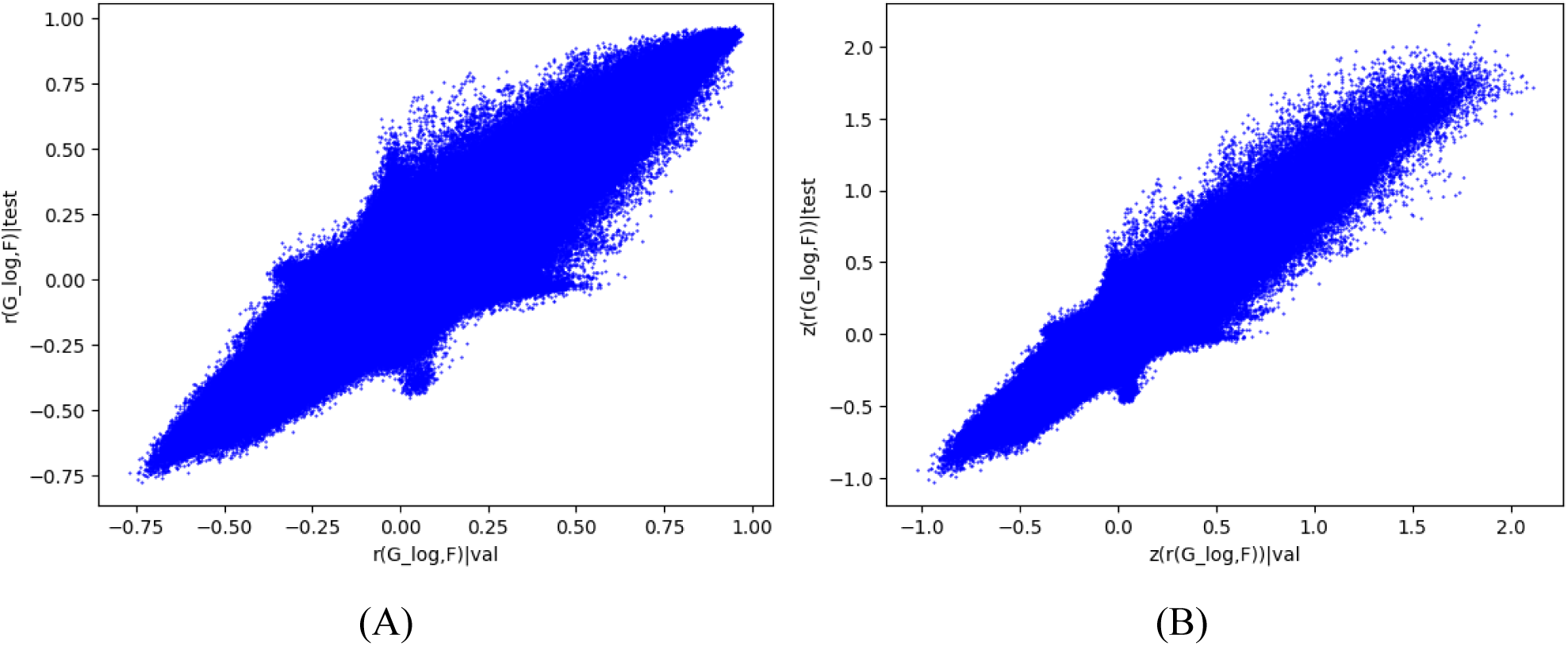
Reproducibility of gene-feature correlations between datasets. Scatter plots of gene-feature correlations computed on the validation- and respectively test dataset. (A) scatter plot of *correlations* (*R*=0.9205). (B) scatter plot of *Fisher transformed* correlations (*R*=0.9233)

**S1 Table.** The numbers of slides and respectively tiles for each data set and tissue type.

| Tissue | slides |  |  |  | tiles |  |  |  |
| --- | --- | --- | --- | --- | --- | --- | --- | --- |
|  | *train* | *validation* | *test* | *total* | *train* | *validation* | *test* | *total* |
| Adipose - Subcutaneous | 22 | 7 | 7 | 36 | 6,915 | 2,203 | 2,318 | 11,436 |
| Adipose - Visceral (Omentum) | 12 | 5 | 4 | 21 | 3,374 | 1,668 | 1,008 | 6,050 |
| Adrenal Gland | 29 | 9 | 9 | 47 | 10,007 | 3,382 | 2,040 | 15,429 |
| Artery - Aorta | 25 | 8 | 9 | 42 | 7,346 | 2,203 | 2,646 | 12,195 |
| Artery - Coronary | 30 | 10 | 10 | 50 | 1,559 | 816 | 703 | 3,078 |
| Artery - Tibial | 27 | 9 | 9 | 45 | 1,581 | 687 | 529 | 2,797 |
| Bladder | 7 | 2 | 2 | 11 | 4,472 | 1,066 | 1,276 | 6,814 |
| Brain - Cerebellum | 29 | 10 | 10 | 49 | 7,449 | 2,848 | 1,958 | 12,255 |
| Brain - Cortex | 30 | 9 | 10 | 49 | 8,793 | 2,495 | 2,845 | 14,133 |
| Breast - Mammary Tissue | 25 | 9 | 8 | 42 | 7,417 | 3,739 | 3,625 | 14,781 |
| Cervix - Ectocervix | 3 | 2 | 1 | 6 | 1,161 | 908 | 823 | 2,892 |
| Cervix - Endocervix | 3 | 1 | 1 | 5 | 1,375 | 221 | 597 | 2,193 |
| Colon - Sigmoid | 30 | 10 | 10 | 50 | 12,413 | 3,078 | 4,067 | 19,558 |
| Colon - Transverse | 28 | 9 | 10 | 47 | 9,754 | 3,698 | 3,159 | 16,611 |
| Esophagus - Gastroesophageal Junction | 28 | 10 | 10 | 48 | 13,195 | 5,032 | 5,800 | 24,027 |
| Esophagus - Mucosa | 28 | 10 | 10 | 48 | 6,225 | 2,086 | 1,937 | 10,248 |
| Esophagus - Muscularis | 28 | 8 | 9 | 45 | 12,076 | 3,246 | 3,612 | 18,934 |
| Fallopian Tube | 4 | 1 | 2 | 7 | 638 | 214 | 170 | 1,022 |
| Heart - Atrial Appendage | 30 | 10 | 10 | 50 | 10,691 | 3,833 | 4,337 | 18,861 |
| Heart - Left Ventricle | 29 | 9 | 9 | 47 | 10,861 | 3,021 | 2,772 | 16,654 |
| Kidney - Cortex | 27 | 9 | 9 | 45 | 10,139 | 3,227 | 3,365 | 16,731 |
| Liver | 30 | 10 | 10 | 50 | 13,229 | 4,111 | 4,197 | 21,537 |
| Lung | 30 | 10 | 9 | 49 | 10,314 | 3,181 | 3,278 | 16,773 |
| Minor Salivary Gland | 30 | 10 | 10 | 50 | 4,426 | 1,527 | 1,878 | 7,831 |
| Muscle - Skeletal | 30 | 10 | 10 | 50 | 12,499 | 3,732 | 4,059 | 20,290 |
| Nerve - Tibial | 29 | 9 | 9 | 47 | 3,359 | 1,365 | 789 | 5,513 |
| Ovary | 29 | 10 | 9 | 48 | 12,178 | 3,475 | 3,594 | 19,247 |
| Uterus | 29 | 10 | 10 | 49 | 12,229 | 4,281 | 4,025 | 20,535 |
| Vagina | 30 | 10 | 10 | 50 | 16,274 | 5,171 | 5,312 | 26,757 |
| Pancreas | 29 | 9 | 10 | 48 | 12,863 | 3,965 | 4,799 | 21,627 |
| Pituitary | 30 | 10 | 9 | 49 | 4,940 | 1,598 | 1,793 | 8,331 |
| Prostate | 29 | 9 | 10 | 48 | 13,939 | 3,718 | 4,792 | 22,449 |
| Skin - Not Sun Exposed | 30 | 10 | 10 | 50 | 13,990 | 4,644 | 4,128 | 22,762 |
| Skin - Sun Exposed | 30 | 10 | 10 | 50 | 11,582 | 4,911 | 3,545 | 20,038 |
| Small Intestine - Terminal Ileum | 29 | 10 | 10 | 49 | 10,455 | 3,158 | 3,375 | 16,988 |
| Spleen | 30 | 9 | 10 | 49 | 12,747 | 4,595 | 4,530 | 21,872 |
| Stomach | 30 | 10 | 9 | 49 | 16,753 | 4,689 | 3,799 | 25,241 |
| Testis | 30 | 8 | 10 | 48 | 9,556 | 2,891 | 3,627 | 16,074 |
| Thyroid | 28 | 9 | 10 | 47 | 10,566 | 3,440 | 4,918 | 18,924 |
| Total | 1,006 | 330 | 334 | 1,670 | 349,340 | 114,123 | 116,025 | 579,488 |

**S2 Table.**
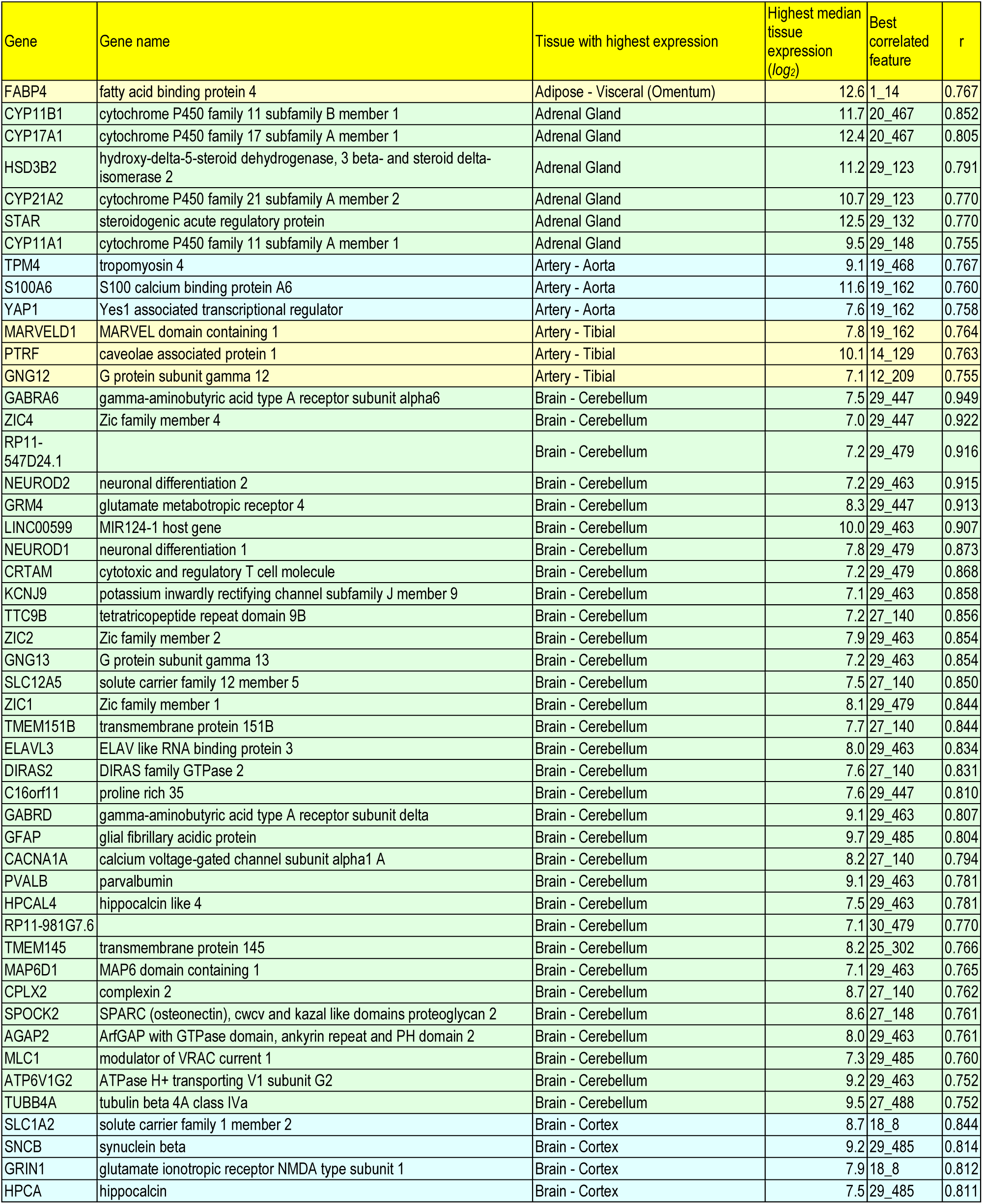
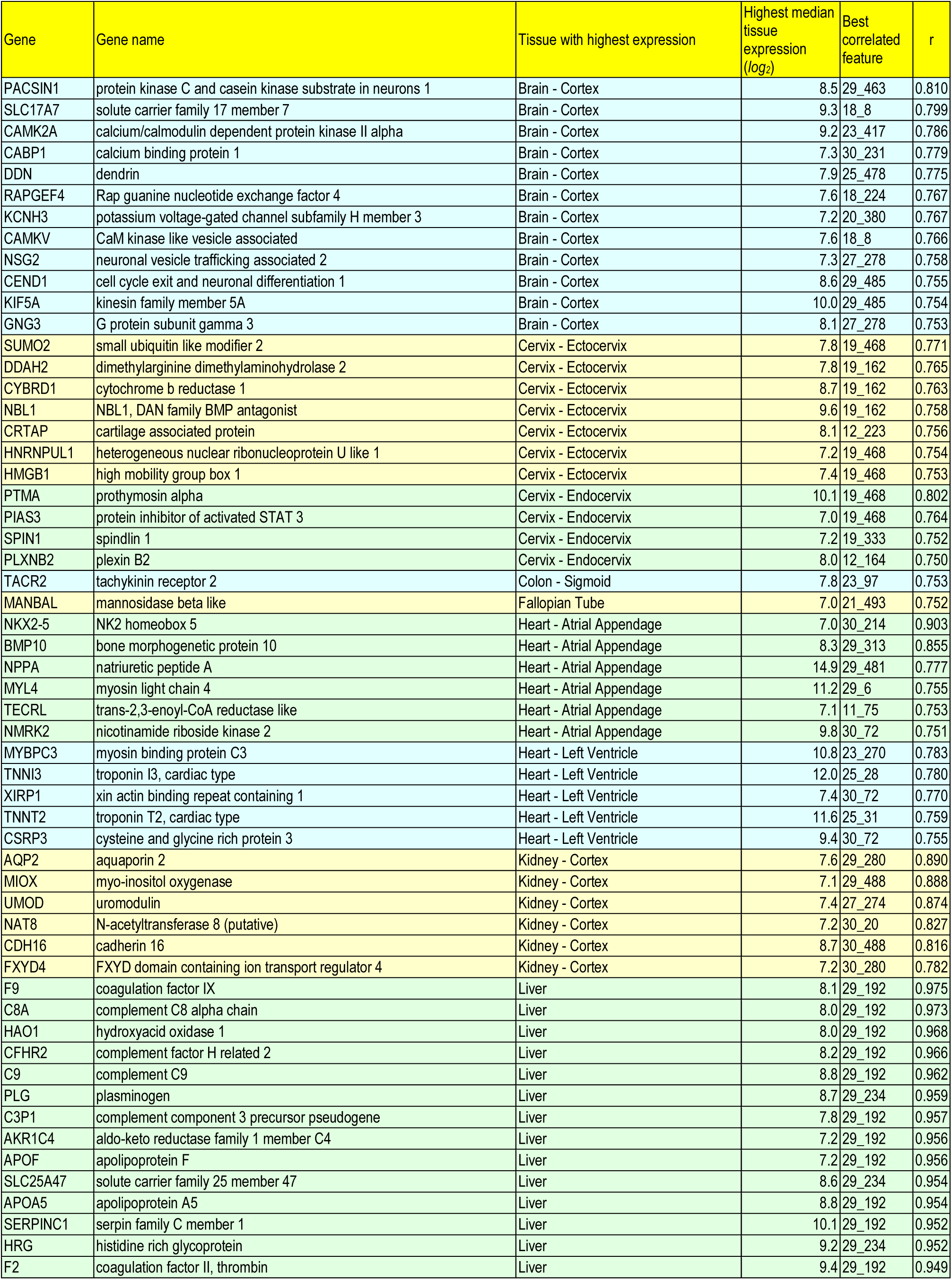
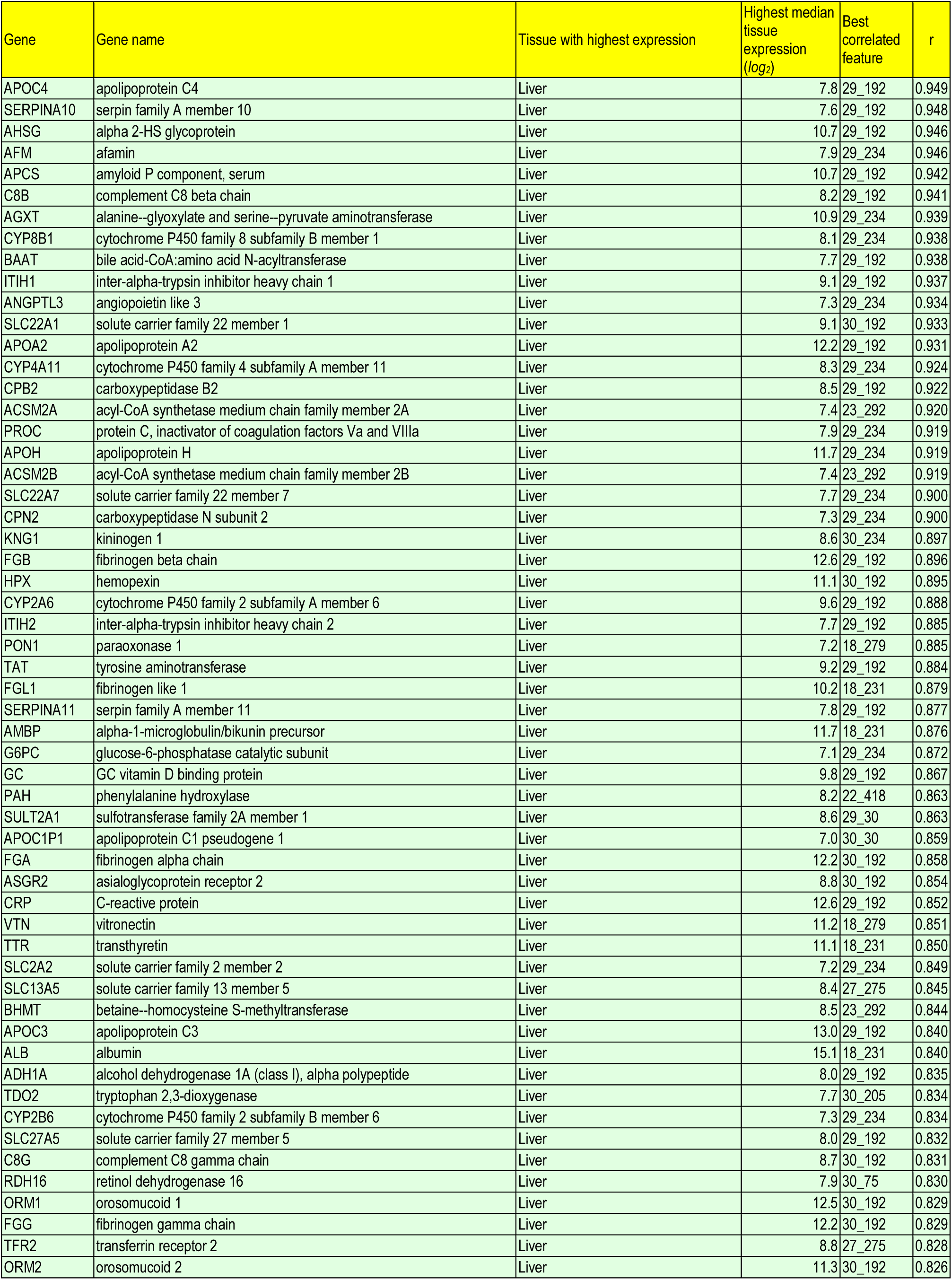
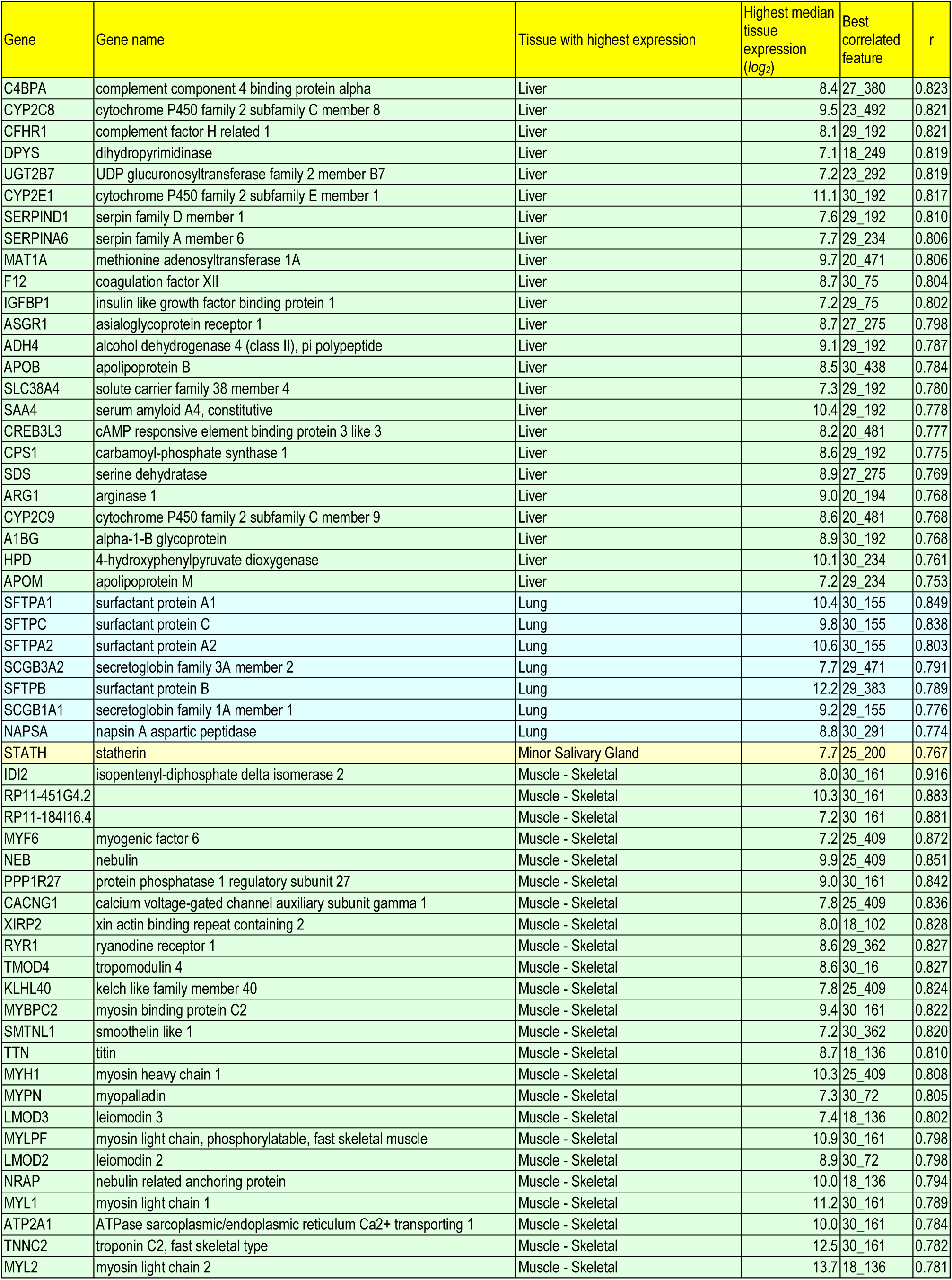
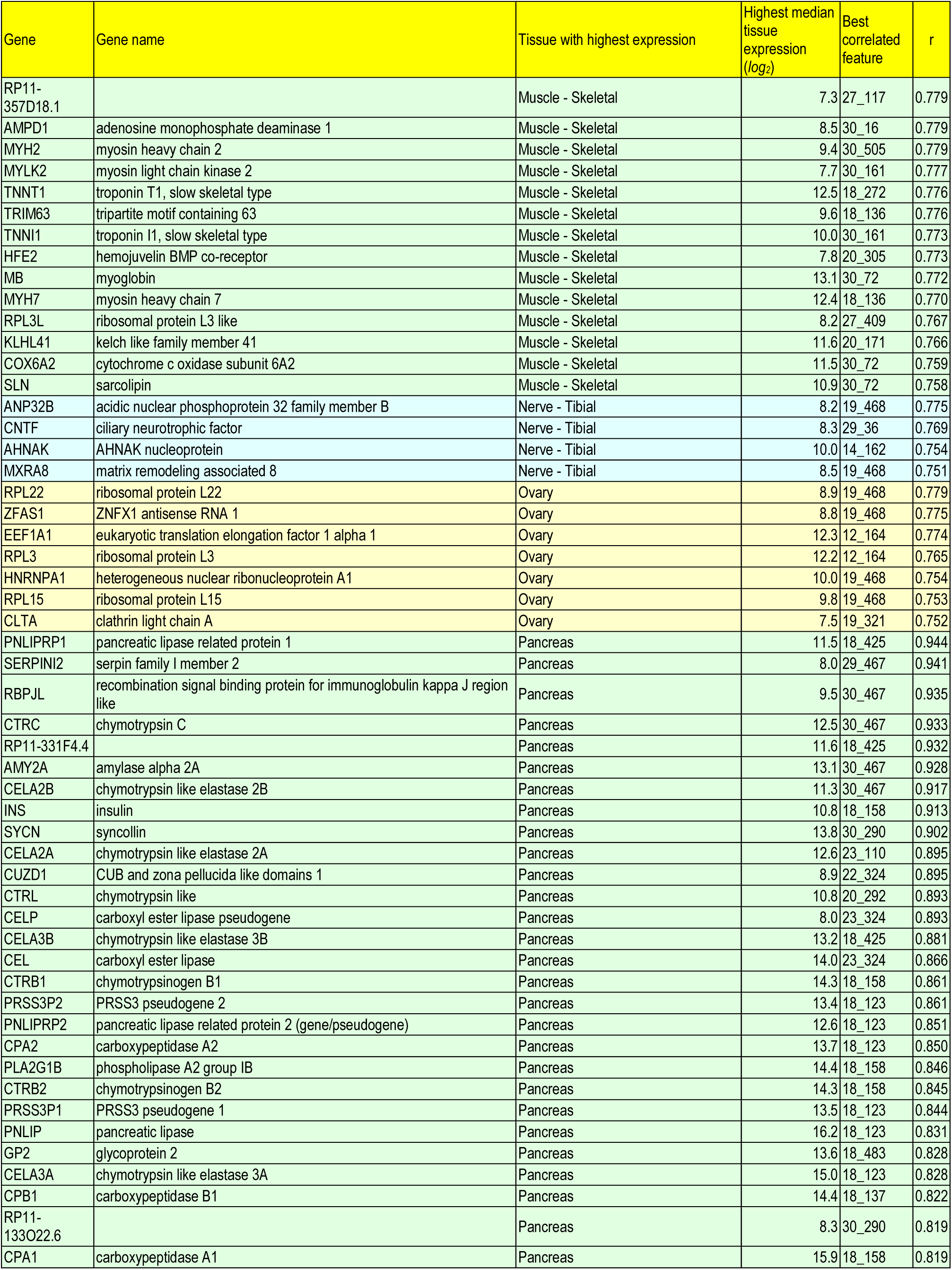
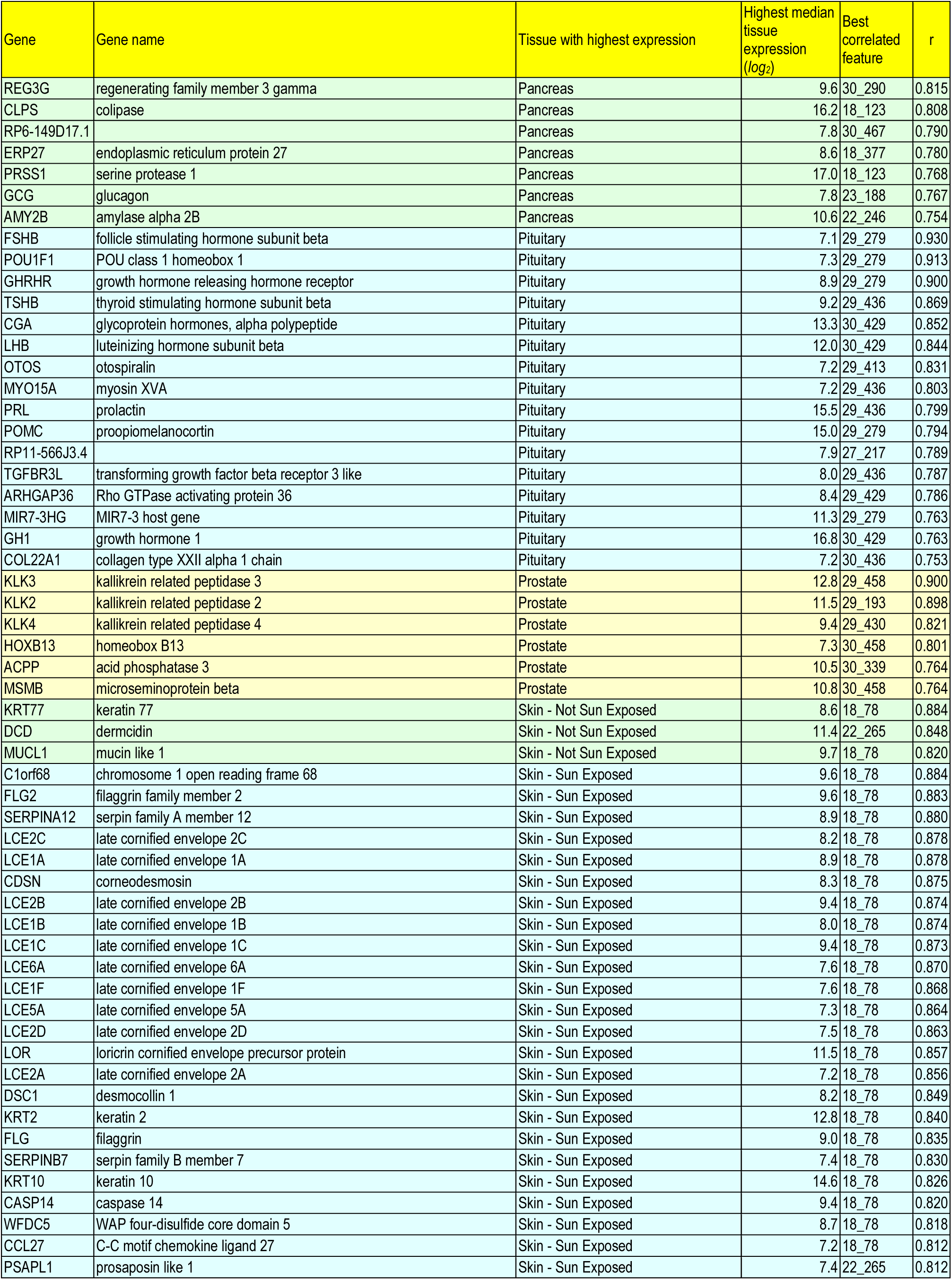
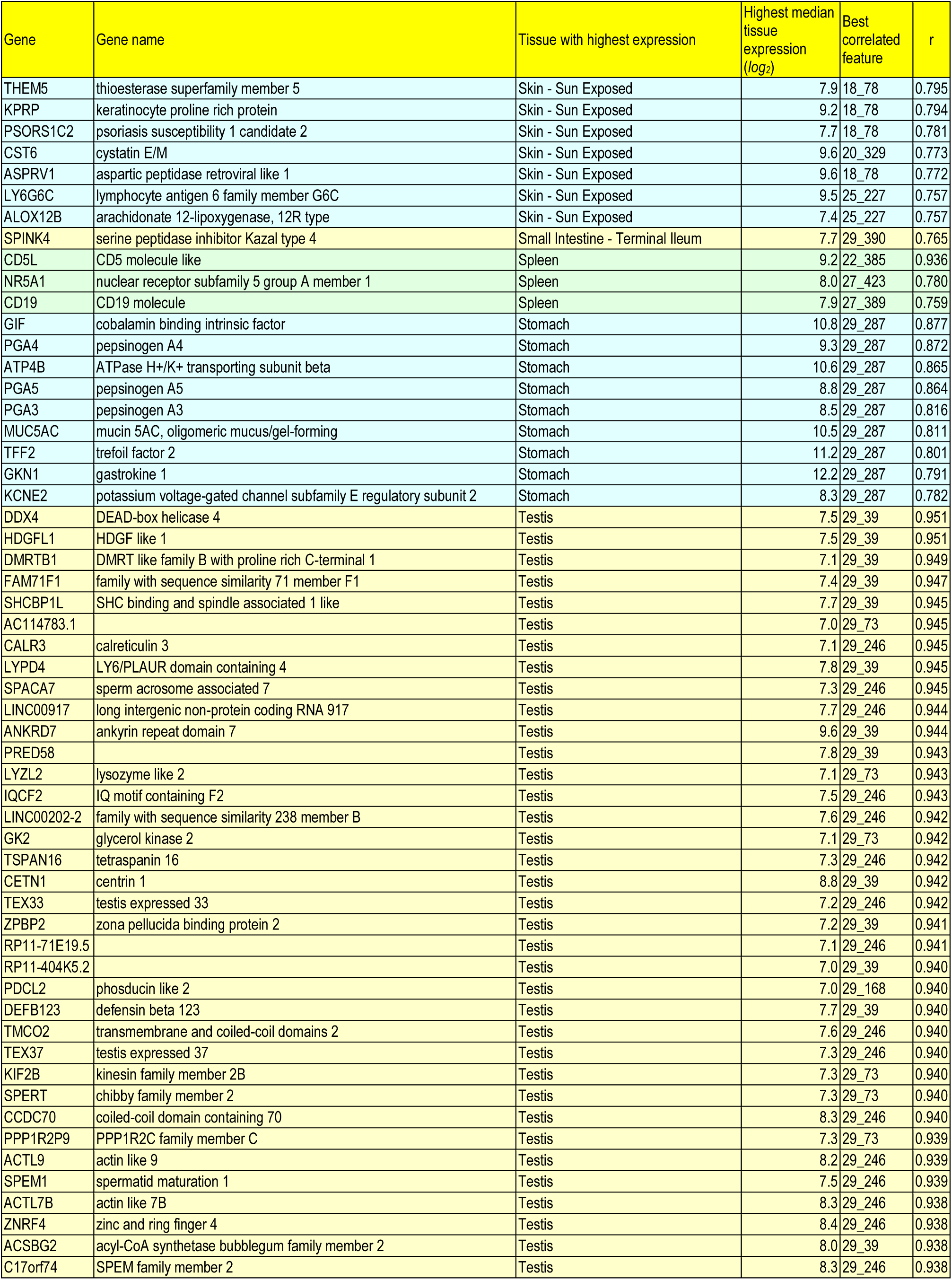
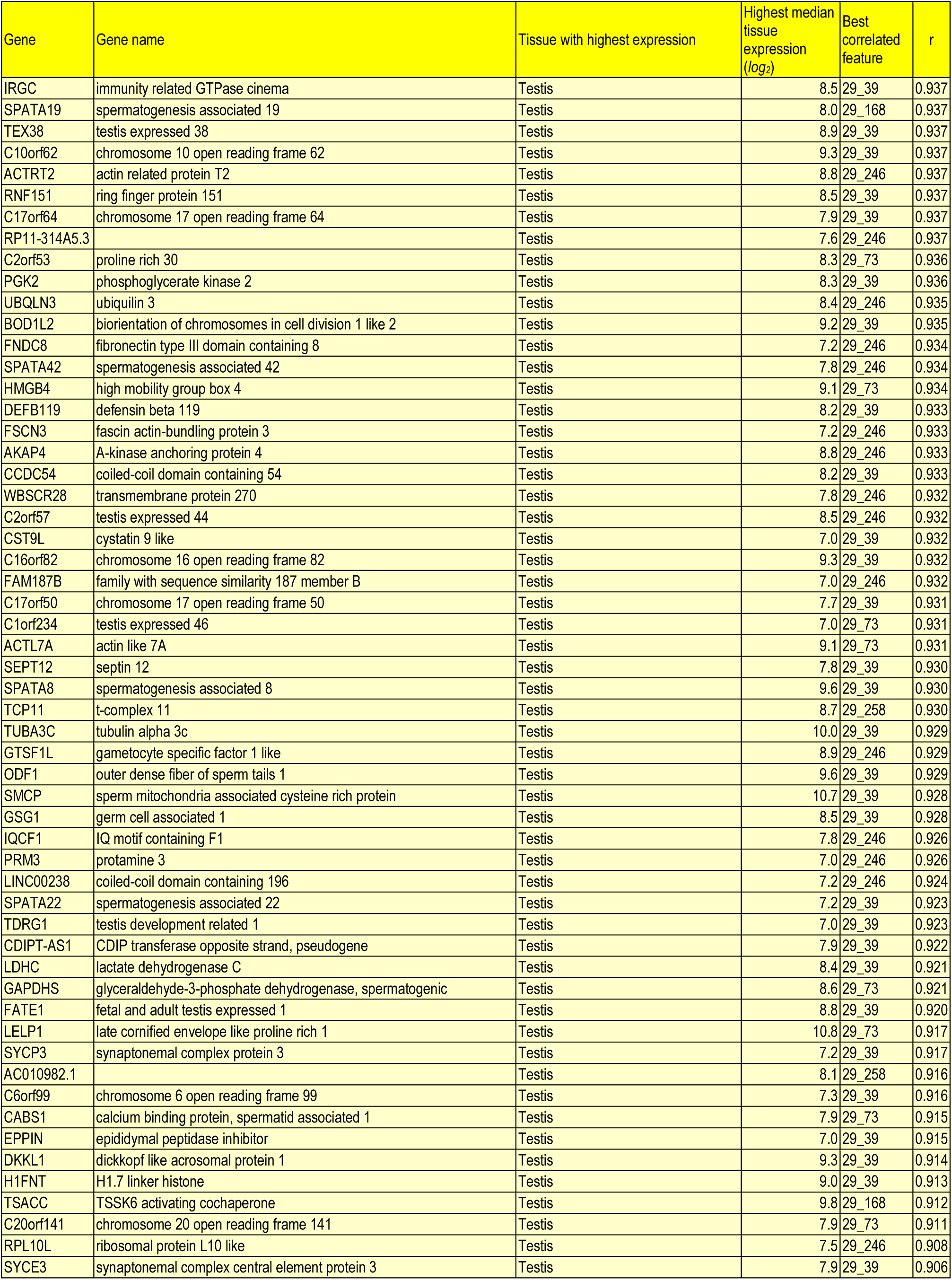
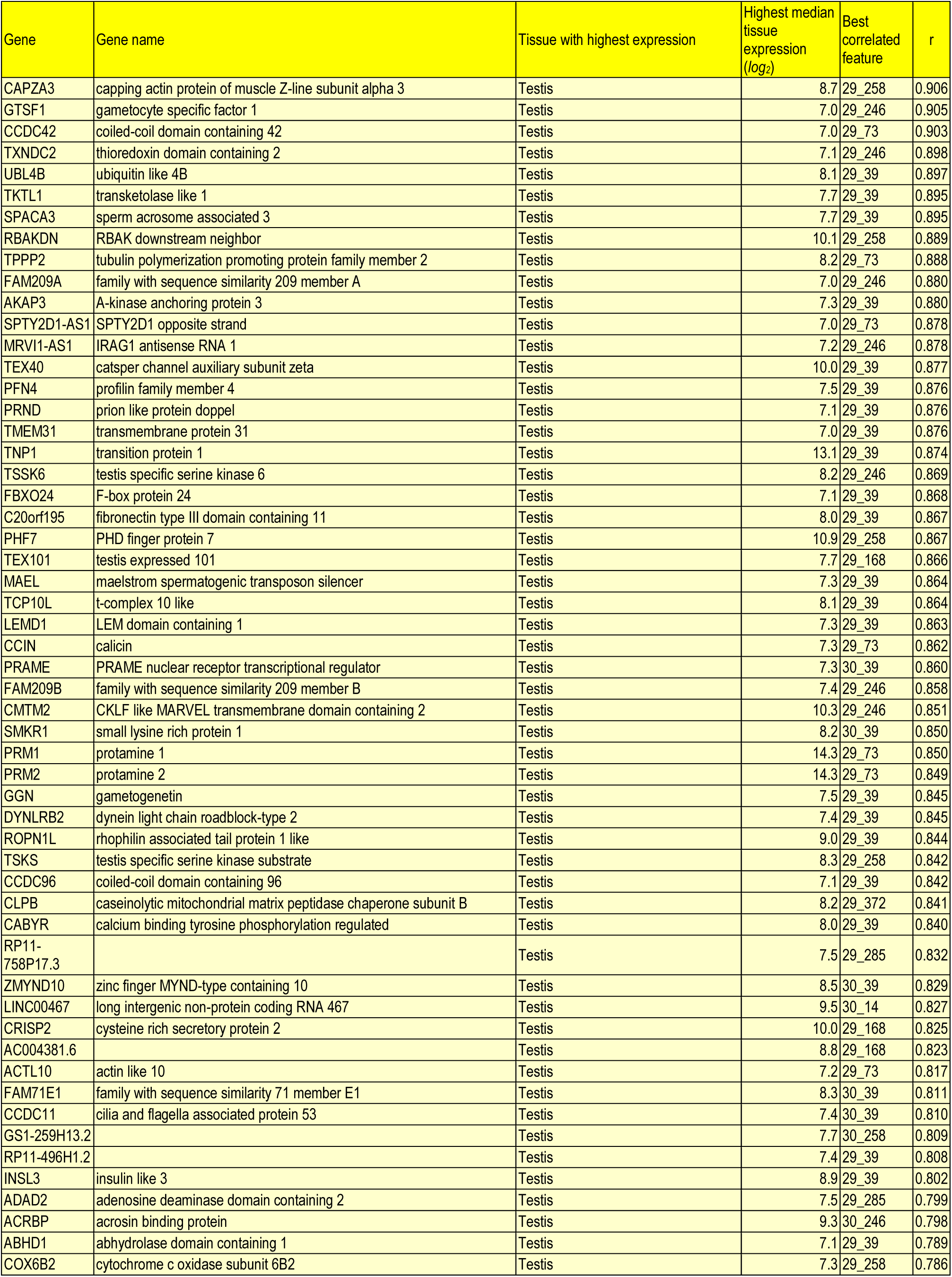
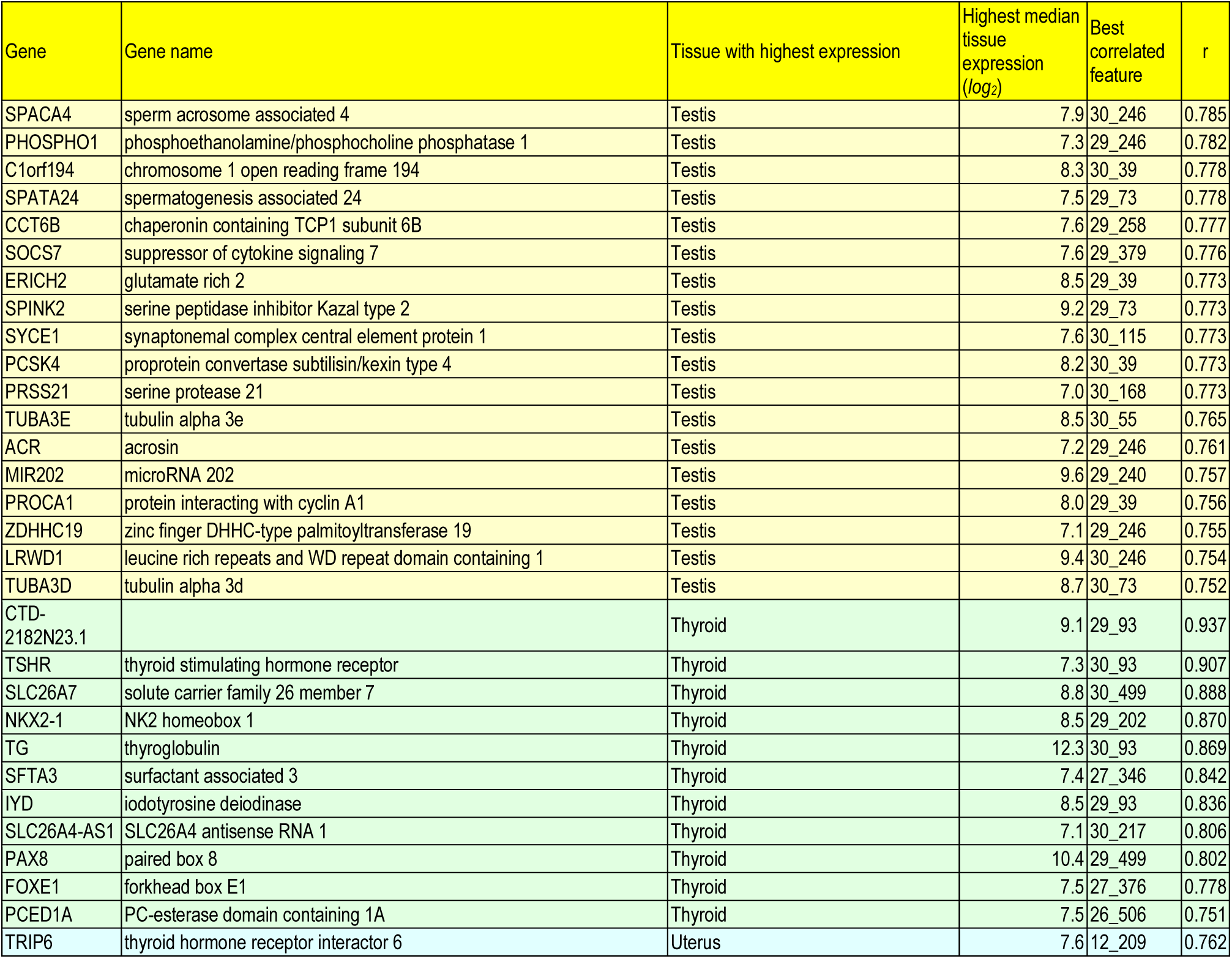
Genes correlated with visual histopathological features. Genes are grouped w.r.t. tissues and ordered by correlation - for each gene we only show the best correlated feature, in the form [layer]_[channel] (for the architecture VGG16 and the test dataset; correlation threshold=0.75, *log*_*2*_ gene expression threshold=7).

**S3 Table.**
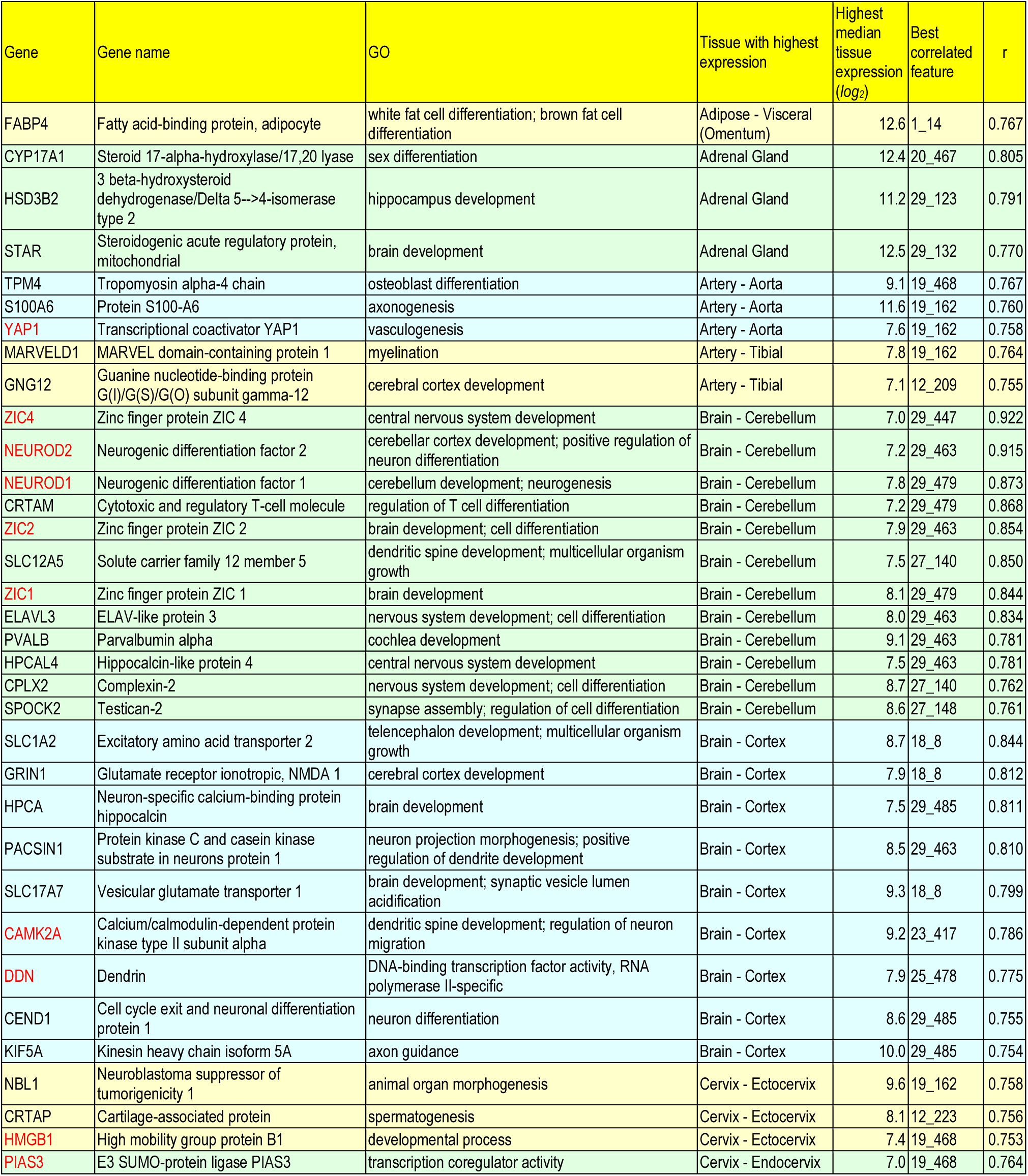
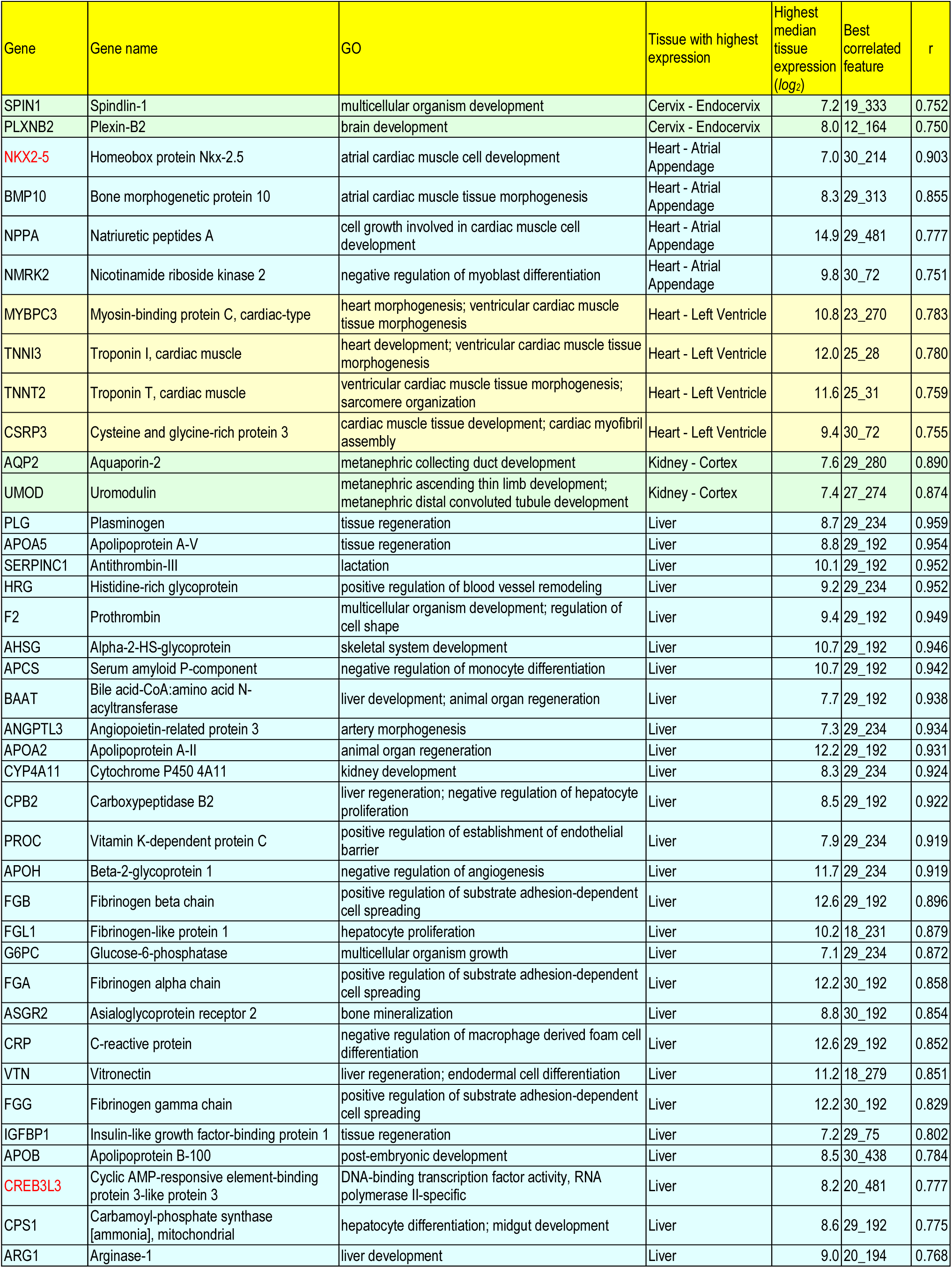
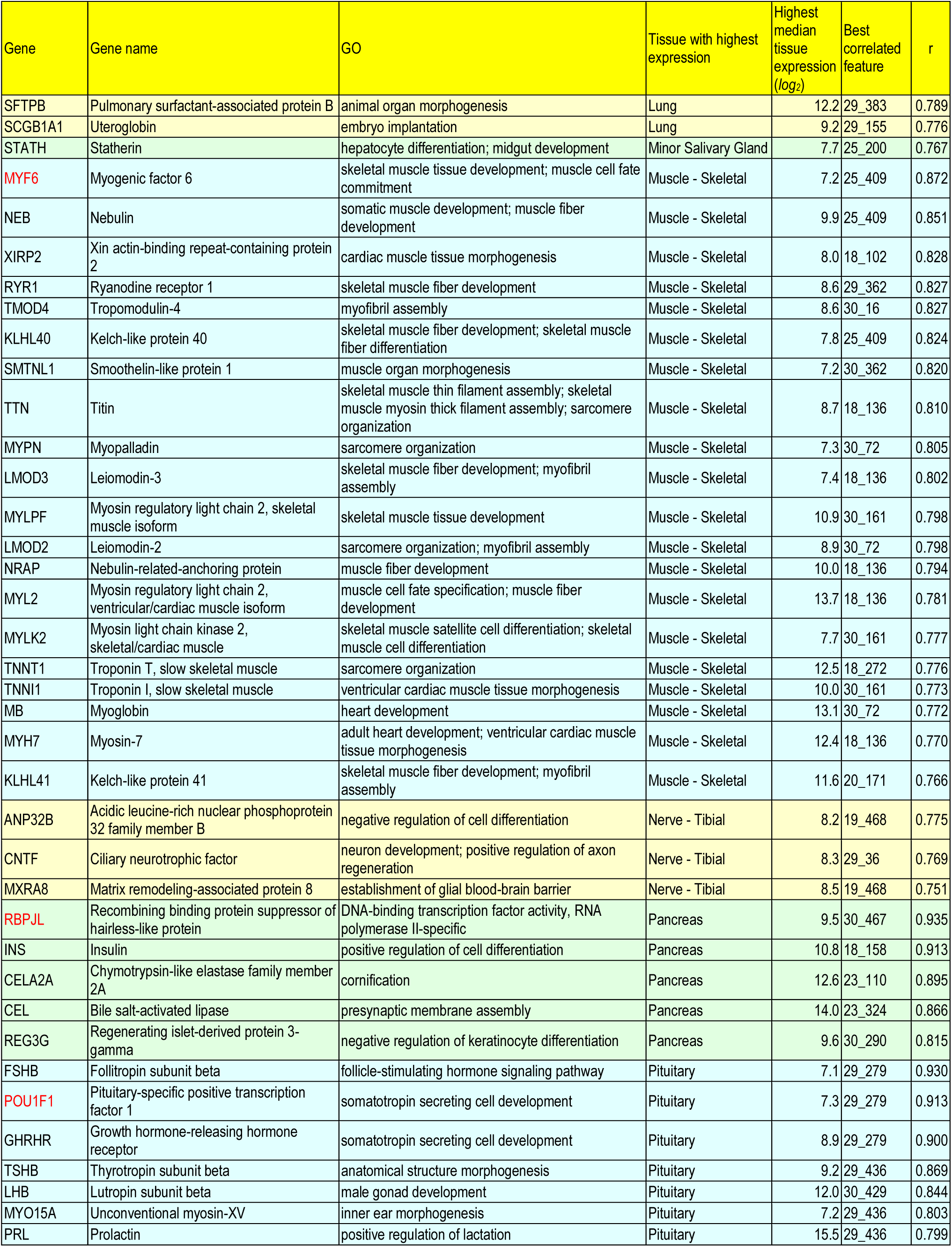
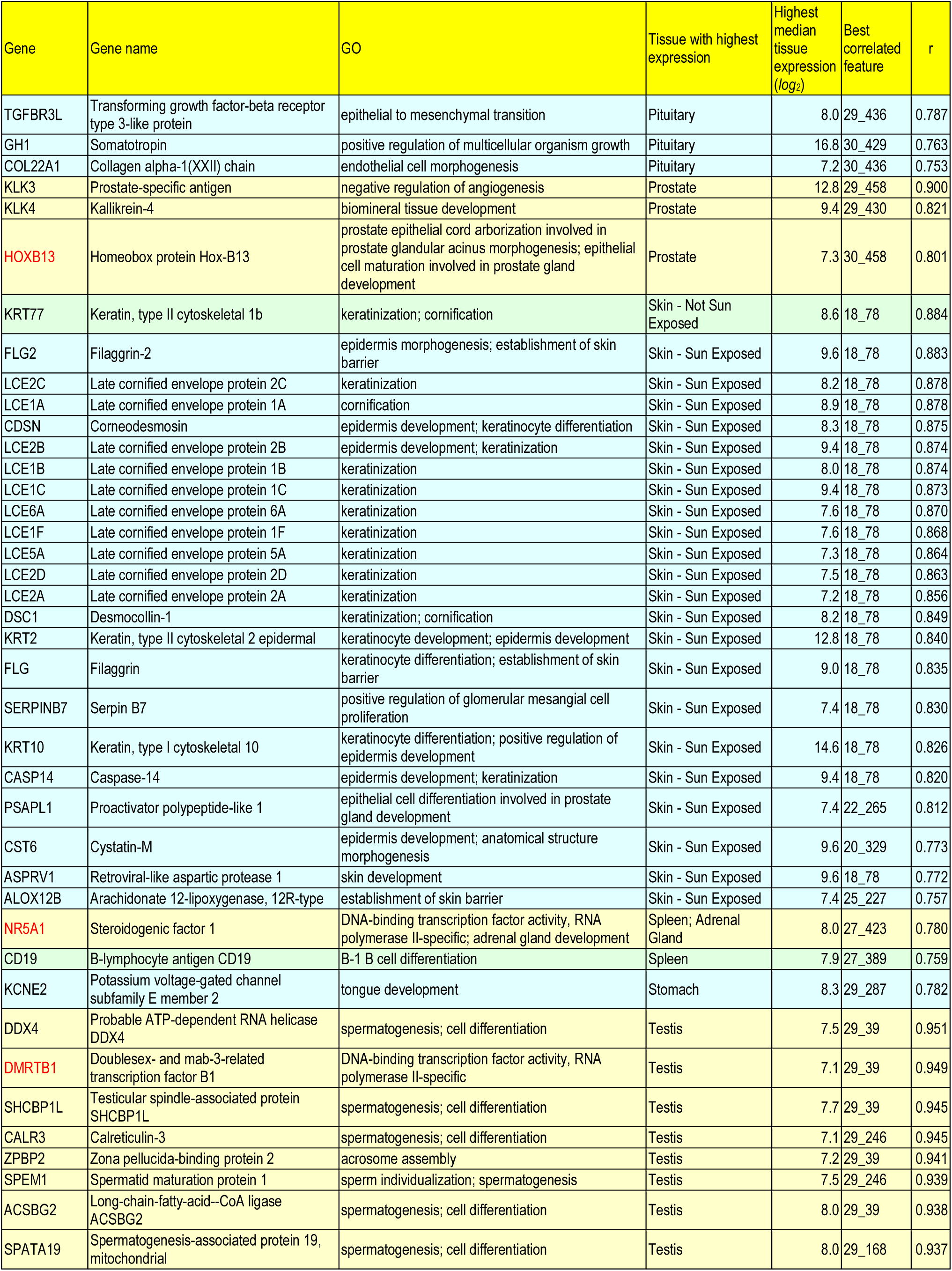
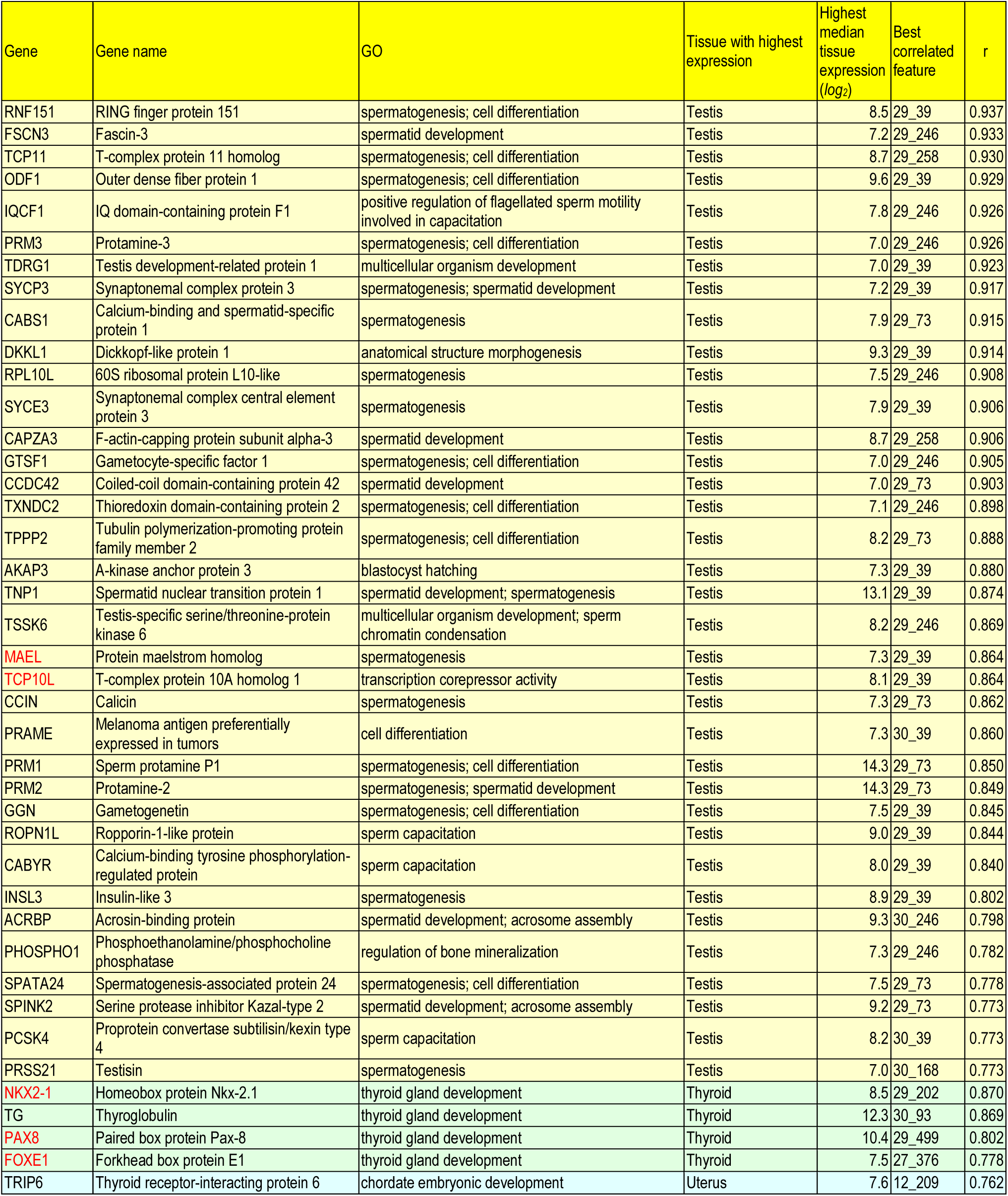
Developmental and transcription regulation genes correlated with visual features. Genes with Gene Ontology ‘*developmental process*’ or ‘*transcription regulator activity*’ annotations are grouped w.r.t. tissues and ordered by correlation - for each gene we only show the best correlated feature, in the form [layer]_[channel] (for the architecture VGG16 and the test dataset; correlation threshold=0.75, *log*_*2*_ gene expression threshold=7). Genes with ‘*transcription regulator activity*’ are shown in red.

**S4 Table.**
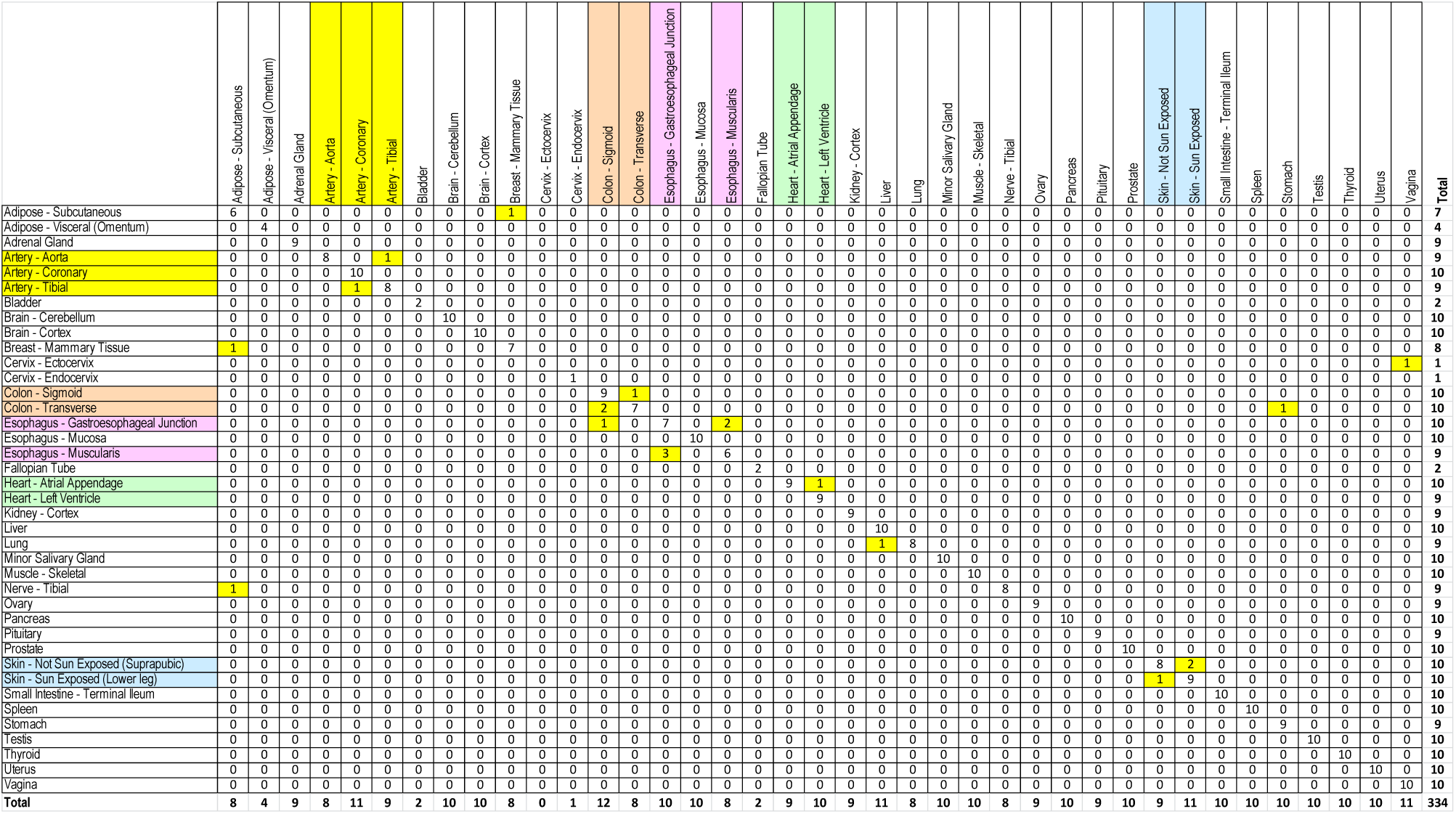
Confusion matrix for the whole slide tissue classifier. VGG16 architecture, test dataset. The rows correspond to the ground truth, the columns to the predicted classes. Similar tissues have been marked with colors, e.g. (Artery - Aorta, Artery - Coronary, Artery - Tibial).

## S1 File. Supporting information

### Selection of representative whole slide image tiles

Unfortunately, guided backpropagation visualization cannot be applied exhaustively because it would involve an unmanageably large number of feature-histological image (*f,X*) pairs. To deal with this problem, we developed an algorithm for selecting a small number of representative histological image tiles to be visualized with guided backpropagation.

**select *k*=3 representative samples for each tissue:**

consider the (gene, feature, tissue) table (*g,f,t*) with corr(*g,f*) ≥ 0.8 and highest median tissue

*log*_*2*_ gene expression ≥ 10

for each tissue *t*

get the samples of the given tissue: samples(*t*)

get the features *f* such that (*f,t*) appear in the table

construct the matrix val(*f,s*) = the value of the feature *f* in sample *s*, for *s* in samples(*t*) and *f* such that (*f,t*) in table

normalize the rows of this matrix: val(*f,s*) = val(*f,s*) / norm_*s’*_ val(*f,s’*)

select the samples with the *k* largest values of min_*f*_ val(*f,s*) as representative samples for tissue *t*: selected_samples(*t*) = *k-*argmax_*s*_ min_*f*_ val(*f,s*)

Note that min_*f*_ val(*f,s*) ensures that if the sample *s* is chosen as representative for tissue *t*, then all features *f* associated to this tissue have a value at least min_*f*_ val(*f,s*).

The same algorithm is applied for selecting a single representative tile from each sample.

### Guided backpropagation image normalization

Guided backpropagation generates images of gradients *X* that must be normalized for visualization. We use the following *log*_*2*_ transformation of the gradients for better emphasizing the lower intensity details:

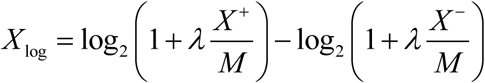

Where 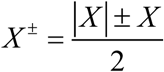, *M* = max(max(*X* ^+^), max(*X* ^−^)), *λ*=10. In turn, *X*_log_ is further normalized to the range [0,1]:

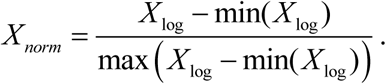

